# eEF1A2 controls local translation and actin dynamics in structural synaptic plasticity

**DOI:** 10.1101/2020.10.20.346858

**Authors:** Mònica B. Mendoza, Sara Gutierrez, Raúl Ortiz, David F. Moreno, Maria Dermit, Martin Dodel, Elena Rebollo, Miquel Bosch, Faraz K. Mardakheh, Carme Gallego

## Abstract

Synaptic plasticity involves structural modifications in dendritic spines. Increasing evidence suggests that structural plasticity is modulated by local protein synthesis and actin remodeling in a synapsis-specific manner. However, the precise molecular mechanisms connecting synaptic stimulation to these processes in dendritic spines are still unclear. In the present study, we demonstrate that the configuration of phosphorylation sites in eEF1A2, an essential translation elongation factor in neurons, is a key modulator of structural plasticity in dendritic spines. A mutant that cannot be phosphorylated stimulates translation but reduces actin dynamics and spine density. By contrast, the phosphomimetic variant loosens its association with F-actin and becomes inactive as a translation elongation factor. Metabotropic glutamate receptor signaling triggers a transient dissociation of eEF1A2 from its GEF protein in dendritic spines, in a phospho-dependent manner. We propose that eEF1A2 establishes a crosstalk mechanism that coordinates local translation and actin dynamics during spine remodeling.

## Introduction

Dendritic spines mediate the vast majority of excitatory synaptic transmission events in the mammalian brain. Structural changes in dendritic spines are essential for synaptic plasticity and brain development (Holtmaat and Svoboda, 2009). The total excitatory input that a neuron can receive is dependent on the complexity of the dendritic network and the density and morphology of spines. Small alterations in average spine density and size may reveal a profound dysfunction at the cellular or circuit level (Forrest et al., 2018). Inside dendritic spines, biochemical states and protein-protein interactions are dynamically modulated by synaptic activity, leading to the regulation of protein synthesis and reorganization of the actin cytoskeleton (Nakahata and Yasuda, 2018). An increasing number of studies support the idea that the actin cytoskeleton and the translation machinery are intrinsically connected and may show reciprocal regulation (DeRubeis et al., 2013; Huang et al., 2013; Santini et al., 2017). Perturbation of the actin cytoskeleton is associated with a dramatic reduction in the rate of global protein synthesis in yeast and mammalian cells (Kim and Coulombe, 2010; Piper et al., 2015).

Regulation of mRNA translation initiation and elongation is essential for synaptic plasticity and memory formation (Gal-Ben-Ari et al., 2012; Kapur et al., 2017; Sossin and Costa-Mattioli, 2019). Studies on the regulation of translation have traditionally focused on the initiation step. There is, however, growing evidence that the elongation step is also regulated to achieve a more robust transient control of the translational machinery in response to synaptic activity(Kenney et al., 2014; Wang et al., 2010). The eukaryotic elongation factor 1 alpha (eEF1A) is a classic G-protein that delivers aminoacylated tRNAs to the A site of the ribosome during translation elongation in a GTP-dependent manner. Recycling of the inactive eEF1A-GDP complex back to the active GTP-bound state is mediated by the eEF1B complex, which acts as a guanine nucleotide exchange factor (GEF) (Negrutskii and El’skaya, 1998). In addition to its well established function in protein synthesis, a number of non-canonical functions have been reported for eEF1A (Sasikumar et al., 2012). The most studied of these is the ability of eEF1A to interact with and modulate the actin cytoskeleton (Bunai et al., 2006; Gross and Kinzy, 2005; Liu et al., 1996).

Vertebrates have two *eEF1A* genes that encode different isoforms, eEF1A1 and eEF1A2. Intriguingly, these isoforms are 92% identical at the amino acid level (Soares et al., 2009) and yet they display very different expression patterns. Isoform eEF1A1 is expressed ubiquitously during development but is replaced by isoform eEF1A2 in neurons and muscle cells over the course of postnatal development (Khalyfa et al., 2001). This expression switch is a vital process, and the complete loss of function of the isoform eEF1A2 in the mouse causes severe neurodegeneration, loss of muscle bulk and death by 4 weeks (Chambers et al., 1998). Despite the fact that numerous studies have been published on the two eEF1A variants, the reasons underlying the developmental switch between the two eEF1A isoforms in neurons and muscle cells remain poorly understood.

eEF1A displays a large repertoire of post-translational modifications brought about by phosphorylation, most of them occurring within conserved regions of both isoforms (Negrutskii et al., 2012; Soares et al., 2009; Soares and Abbott, 2013). As an interesting exception, it has been reported that the kinase receptor for activated protein C kinase 1 (RACK1) recruits stress-activated c-Jun N-terminal kinase (JNK) to polysomes, where it phosphorylates eEF1A2 at Ser205 and Ser358 and promotes degradation of newly synthesized polypeptides by the proteasome. Since Ser358 is evolutionary conserved but not present in isoform eEF1A1, this post-transcriptional regulatory mechanism could constitute a relevant difference in the physiological roles of the two isoforms (Gandin et al., 2013).

So far, although eEF1A2 is the most abundant isoform in mature neurons, most published work has been carried out in a non-neuronal context. It is worth noting that the developmental timeline of synaptic spines and neuronal circuit formation occurs when isoform eEF1A1 is totally replaced by isoform eEF1A2 in neurons. Another concern is that numerous data from experiments in mammalian cells have been analyzed with antibodies that do not distinguish between the two isoforms. Here, we demonstrate that the configuration of phosphorylation sites unique to the eEF1A2 isoform plays a role in dendritic structural plasticity. We show that expression of a phosphoablated mutant in hippocampal neurons results in a significant reduction in dendritic spine density. By contrast, expression of a phosphomimetic mutant impairs its association with F-actin increasing its mobility and F-actin dynamics both in dendrites and synaptic spines. Regarding the canonical function of eEF1A2, we found that the phosphomimetic variant is unable to sustain translation in yeast cells. Moreover, our study reveals that the stimulation of metabotropic glutamate receptor signaling triggers a transient dissociation of eEF1A2 from its GEF protein in a phosphosite-dependent manner. Our findings demonstrate important mechanistic differences between the two eEF1A isoforms and point to the notion that eEF1A2 locally links synaptic inputs to translation and actin remodeling for structural plasticity in neurons via a phospho-dependent regulation.

## Results

### eEF1A2 phosphosite configuration modulates spine growth

First, we wanted to test whether the eEF1A isoform switch can be reproduced *in vitro.* Consistent with previous studies in mouse brain (Khalyfa et al., 2001), we observed that isoform eEF1A2 expression progressively increased in cultures of hippocampal neurons, becoming the main isoform two weeks after cell plating. The progressive increase was observed at both protein and mRNA levels (Figure 1A-C). Although eEF1A1 and eEF1A2 contain 462 and 463 amino acid residues respectively, isoform eEF1A1 migrated slightly faster as deduced by immunoblot analysis with a specific antibody against isoform eEF1A2 (Figure 1A). Whereas eEF1A2 is exclusively expressed in neurons, eEF1A1 is the main isoform in glial cells (Pan et al., 2004), which explains why we observed low levels of eEF1A1 in long-term hippocampal cultures.

**Figure 1.**
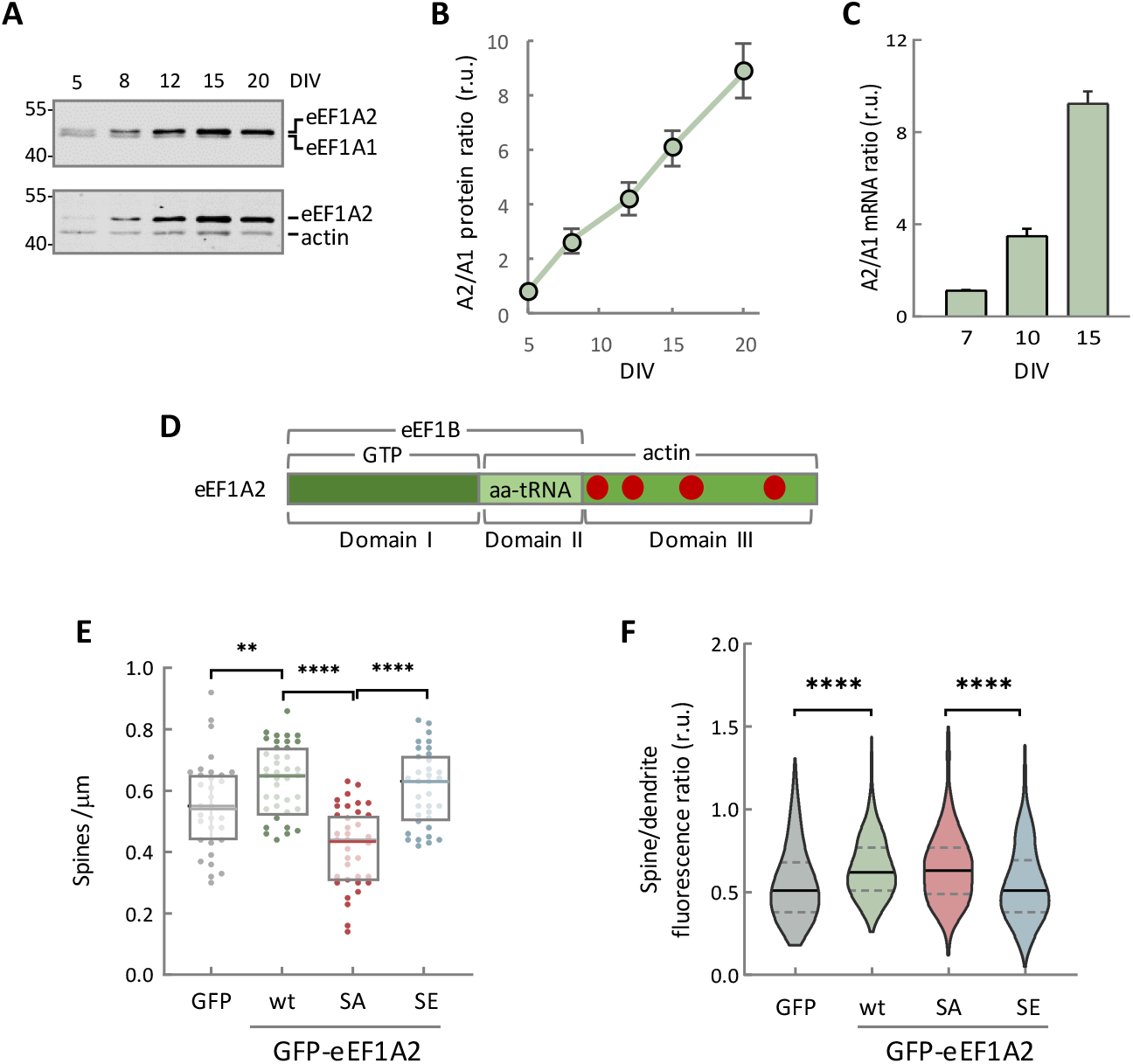
eEF1A2 phosphosite configuration modulates spine growth. (A) Differential expression of eEF1A isoforms in cultured hippocampal neurons from E17 mouse embryos. Samples were collected at different days *in vitro* (DIV) and analyzed by immunoblotting using a mouse monoclonal αeEF1A (upper panel) or a rabbit polyclonal αeEF1A2 (bottom panel). (B) Quantification of eEF1A isoforms from immunoblot analysis as in panel A. eEF1A protein levels were normalized relative to actin. Ratios are represented as mean ± SEM (n=3). (C) Quantification of eEF1A isoform mRNA ratios from cultured hippocampal neurons by qRT-PCR. Bars represent mean ± SEM (n=3). (D) Scheme showing the three domains of eEF1A2 and relevant protein interactions. Red dots represent phosphorylation sites in domain III that are not present in eEF1A1. See Figure 1–figure supplement 1A for full details. (E) Quantification of spine density in CA1 pyramidal neurons. Organotypic hippocampal slices were transfected with plasmids expressing GFP or GFP-fusions of wt, SA or SE eEF1A2 proteins, and a plasmid expressing RFP (DsRed2) for dendritic tracing. Changes in spine number were assessed by analyzing GFP-positive neurons with SpineJ software (Pedraza et al., 2014). The total number of observations (spines/neurons) plotted is as follows: GFP, n=834/32; wt, n=1387/39; SA, n=780/36; SE, n=938/35. Single-neuron data (dots) from three independent experiments and median ± Q values are plotted. The results of Mann-Whitney tests (** p<0.01; **** p<0.0001) are also indicated. (F) Quantification of GFP-fused wt, SA and SE eEF1A2 proteins in spines of CA1 pyramidal neurons. GFP fluorescence in spines was normalized to that in corresponding dendritic shafts. The total number of observations (spines/neurons) plotted is as follows: GFP, n=259/10; wt, n=364/10; SA, n=313/12; SE, n=534/13. Median ± Q values and the results of Mann-Whitney tests (**** p<0.0001) are also shown.

The high resolution structure of the yeast eEF1A factor reveals a compact conformation with three domains displaying multiple mutual interactions (Andersen et al., 2001, 2000). While domain I contains the GTP-binding site, domain II is implicated in the interaction with aminoacyl-tRNA. Both domains interact with eEF1Bα during the exchange of GDP for GTP. Finally, domain II and domain III carry residues important for the interaction of eEF1A with the actin cytoskeleton (Liu et al., 2002; Mateyak and Kinzy, 2010; Soares et al., 2009). In domain III, isoform eEF1A2 presents four putative phosphorylation residues, Ser342, Ser358, Ser393 and Ser445, that are not present in isoform eEF1A1 (Figure 1D and Figure 1–figure supplement 1A). Ser358 is conserved in organisms that only have one eEF1A isoform but is restricted to eEF1A2 once this isoform appears in evolution (Figure 1– supplement 1A). This residue is known to be phosphorylated by polysome-associated JNK in response to DHPG (Gandin et al., 2013).

To test whether phosphorylation in domain III is relevant to eEF1A2 function in synaptic plasticity, we replaced the four eEF1A2-specific serines with alanine or glutamic acid to obtain phosphoablated (SA) and phosphomimetic (SE) mutants, respectively. CA1 pyramidal cells of rat hippocampal slice cultures were co-transfected with GFP or GFP-eEF1A2 proteins (wt, SA and SE) and DsRed2 to analyze spine density. The phosphoablated mutant showed a significant reduction in the number of dendritic spines compared to wt and mutant SE (Figure 1E). These results were also confirmed using dissociated hippocampal neurons (Figure 1–figure supplement 1B). We then estimated eEF1A2 distribution by comparing the GFP signal in spines versus the adjacent dendritic shafts. The SE mutant showed a reduced accumulation in spines compared to the GFP-eEF1A2 wt and SA mutant and similar to the levels of GFP (Figure 1F), suggesting that phosphorylation in domain III modulates eEF1A2 targeting to spines. In all, our data point to the notion that eEF1A2 phosphorylation is important for regulation of structural synaptic plasticity.

### Interactome analysis of eEF1A2 phosphomutants dissects translational and non-canonical functions

To elucidate the role of eEF1A2 phosphorylation we decided to examine the interactomes of both the SA and SE mutants. αFLAG immunoprecipitates from HEK293T cells transfected with *FLAG-eEF1A2^SA^* or *FLAG-eEF1A2^SE^* cDNAs were analyzed by LC-MS/MS. Out of a total of 3026 proteins identified, 37 proteins were differentially enriched in SA immunoprecipitates (SA-IP) and 88 proteins in SE immunoprecipitates (SE-IP) (Appendix 1—table 1). Gene ontology (GO) enrichment analysis showed that proteins associated with ribosome biogenesis and translational elongation were significantly (p<0.001) overrepresented in SA-IP compared to SE-IP (Figures 2A and Figure 2–figure supplement 1). Among translation-associated proteins we found Valyl-tRNA synthetase 1, eEF1D, eEF1B2, Cysteinyl-tRNA synthetase, eEF1G and eEF1A1 (Figure 2B). The more efficient interaction of the SA mutant with eEF1A1 suggests that eEF1A2 phosphorylation modulates its ability to dimerize. By contrast, the SE-IP showed a significant enrichment of interactors involved in vesicle transport (p<0.001), protein modification (p=0.006, mostly protein ubiquitination), stress response (p=0.024) and nuclear functions such as DNA replication (p=0.030) or nuclear import (p=0.026). In addition to these categories SE-IP showed enrichment in a set of proteins other than actin itself, which are involved in actin cytoskeleton dynamics: Shroom3 (p<0.001), Filamin-B (p<0.001), α-actinin-4 (p=0.008), RhoA (p=0.014) and F-actin-capping β (p=0.010), suggesting that eEF1A2 phosphorylation could be involved in modulating actin dynamics (Gross, 2013; Truebestein et al., 2015) (Figure 2B).

**Figure 2.**
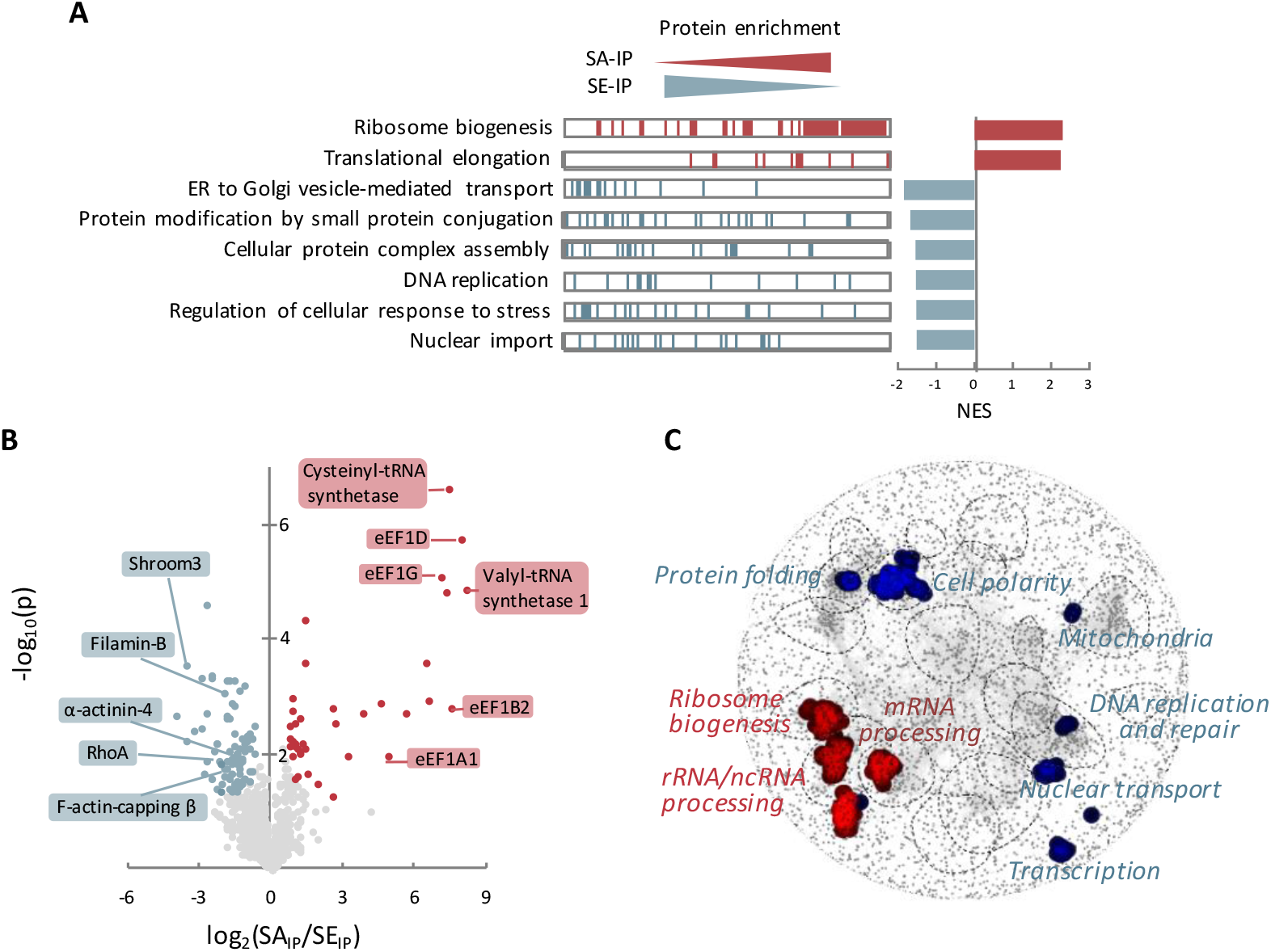
Interactomic analysis of eEF1A2 phosphomutants dissects translational and non-canonical functions. (A) Triplicate immunoprecipitates from HEK293T cells expressing FLAG-tagged SA and SE eEF1A2 proteins were analyzed by LC-MS/MS. The results of a gene set enrichment analysis of FLAG immunoprecipitates (SA-IP and SE-IP) are shown as barcode plots for the most significant GO terms (left). A bar chart with the corresponding normalized enrichment scores (NES) is also shown. See Figure 2–figure supplement 1 and Appendix 1—table 1 for details. (B) Volcano plot showing the relative enrichment of identified interactors in immunoprecipitates of FLAG-tagged SA and SE eEF1A2 proteins. See Appendix 1—table 1 for details. (C) Distribution of eEF1A2 SA (red) and SE (blue) interactor orthologues in the yeast global genetic interaction network. Categories enriched with the corresponding orthologues are indicated.

**Table 1.**
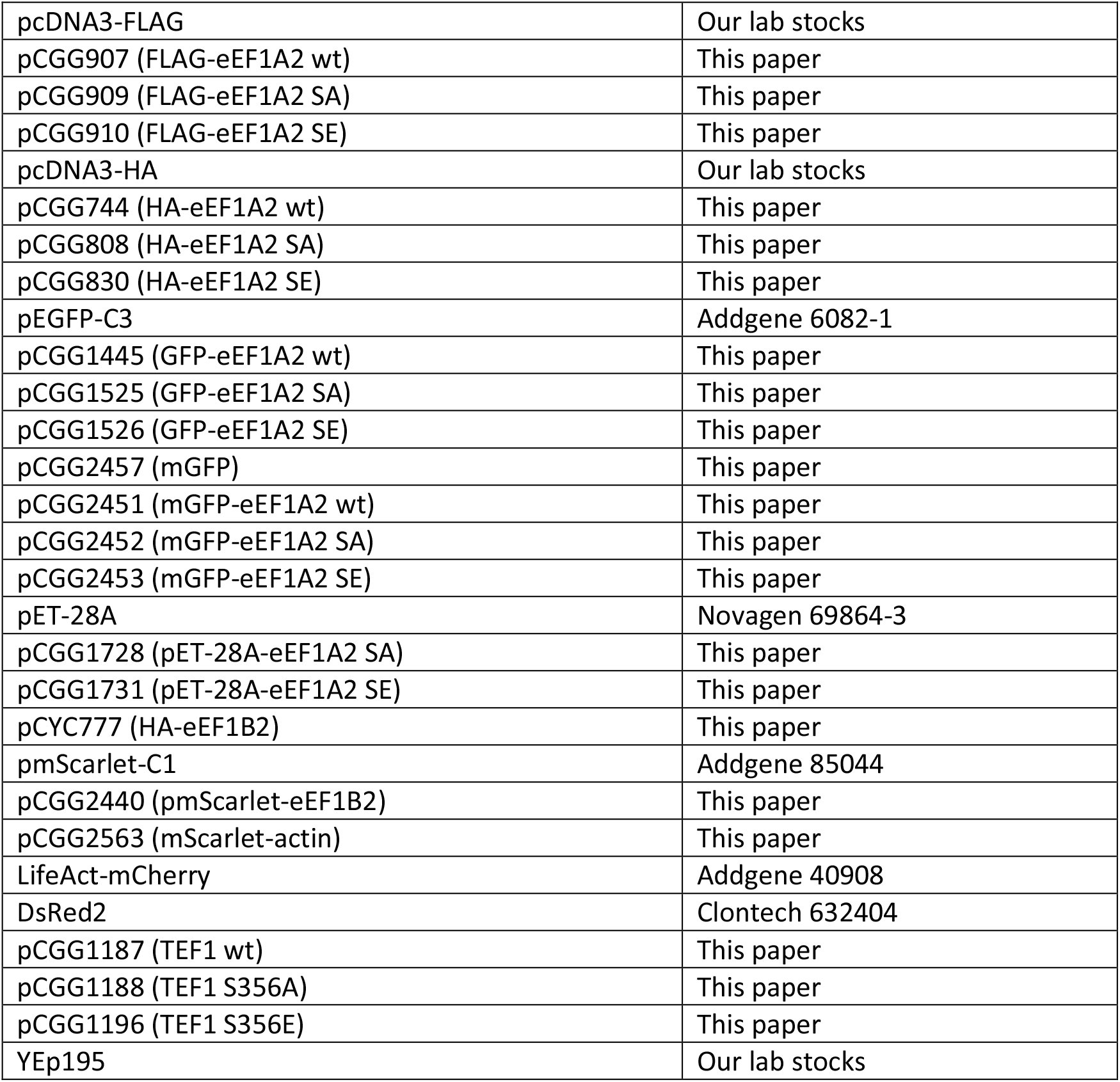
Plasmids used in this study

**Table 2.**
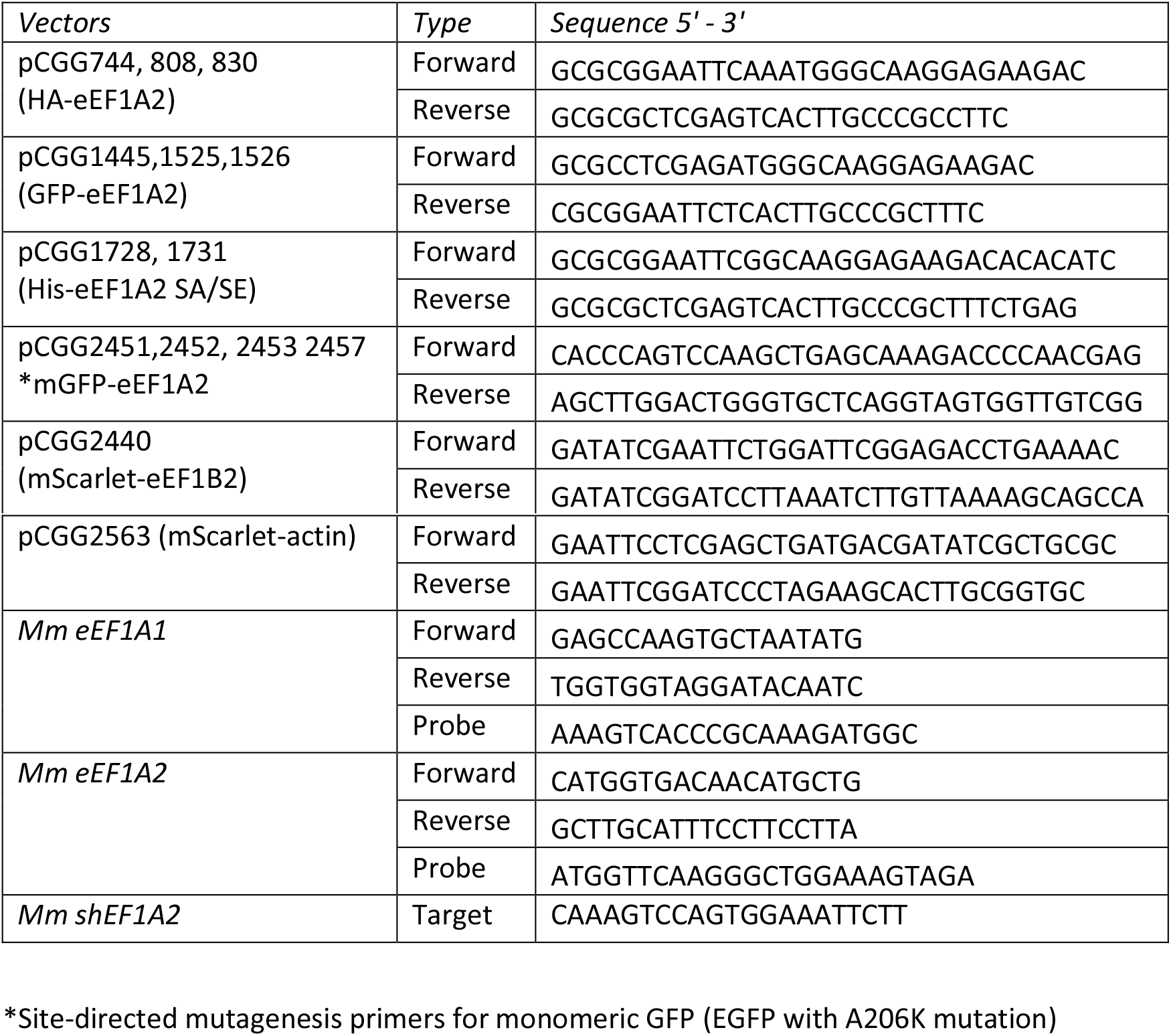
Oligonucleotides used in this study

As a complement to the GO enrichment analysis, we examined how the yeast orthologues of human eEF1A2-phosphomutant interactors are grouped in the global yeast genetic interaction network (Usaj et al., 2017). As shown in Figure 2C, whereas phosphoablated eEF1A2 interactors were found in clusters related to ribosome biogenesis and mRNA processing, phosphomimetic eEF1A2 binding proteins displayed strong genetic interactions in smaller clusters, many of them related to non-canonical functions such as endocytosis, nuclear processes or actin cytoskeleton dynamics.

### Phosphomimetic residues in eEF1A2 hinder its association with F-actin and increase actin dynamics

The first step in remodeling the spine actin network is the unbundling of actin filaments (F-actin), which are normally crosslinked by different types of actin-binding proteins. Dissociation of these actin-crosslinking proteins would allow access to other actin-binding proteins to stimulate spatial-temporal flexibility of the actin filament network (Cingolani and Goda, 2008). To determine whether eEF1A2 phosphorylation could have an effect on its association with F-actin, we used pulldowns (PD) of biotinylated actin to measure the actin-binding properties of phosphomutant-eEF1A2 proteins. The immunoblotting analysis showed that protein levels of the SE mutant bound to actin were remarkably lower compared to wt and SA proteins (Figure 3A and B). In order to further test whether phosphorylation is sufficient to detach eEF1A2 from F-actin, we performed F-actin bundling assays at low speed in which we observed that purified SE mutant completely loses F-actin binding activity (Figure 3C and D).

**Figure 3.**
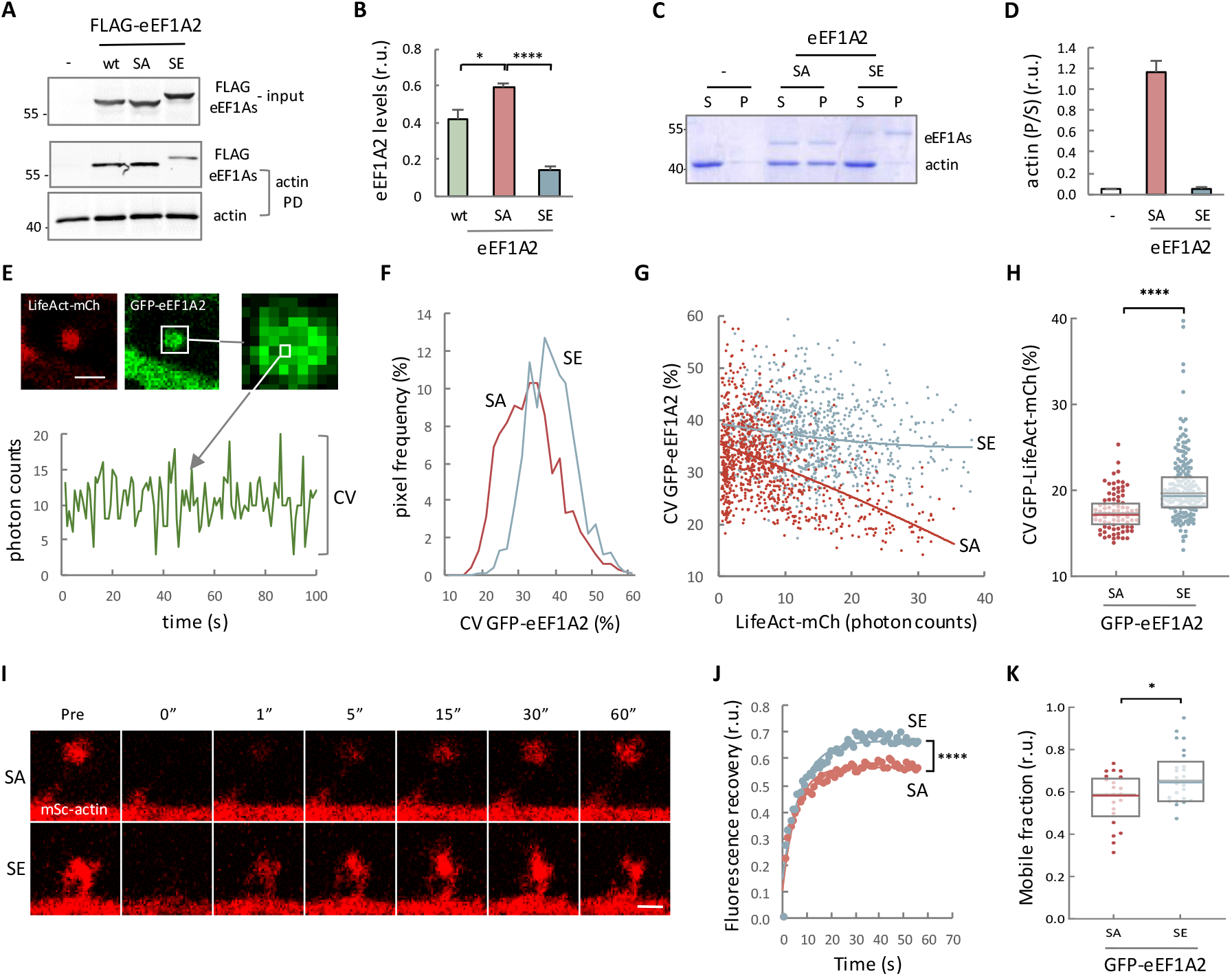
Phosphomimetic residues in eEF1A2 hinder its association with F-actin and increase actin dynamics. (A) Constructs expressing FLAG-tagged eEF1As were transiently transfected into HEK293T cells and, after 24 h, cells lysates were subject to biotin-actin binding assays. Lysate (input) and actin pulldown (PD) samples were analyzed by immunoblotting to detect actin or FLAG-tagged proteins. Empty vector was used as control (-). (B) Quantification of eEF1A isoform ratios from immunoblot analysis as in panel A. Protein levels were normalized relative to input levels. Data are represented as mean ± SEM (n=3), and the results of t tests (* p<0.05, **** p<0.0001) are also shown. (C) Actin bundling assay with His-eEF1A2 fusion proteins. Supernatant (S) and pellet (P) fractions were subjected to SDS-PAGE and stained with Coomassie brilliant blue. (D) Quantification of F-actin levels in pellet and supernatant fractions from actin bundling assays as in panel C. P/S ratios for eEF1A2 mutants (SA and SE) and control (-) are plotted as mean ± SEM (n=3) values. (E) Hippocampal neurons were transfected with plasmids expressing GFP-eEF1A2 and LifeAct-mCh as F-actin reporter, and dendritic spines were analyzed by time-lapse photon-counting microscopy. A representative temporal profile obtained from GFP-eEF1A2 in a single spine pixel is shown at the bottom. Bar, 1 μm. (F) Distributions of the coefficient of variation of fluorescence fluctuations from SA or SE GFP-eEF1A2 proteins as in panel E. The number of observations (pixels/spines) analyzed is as follows: SA, n=1084/12; SE, n=921/16. (G) Coefficient of variation of fluorescence fluctuations from GFP-fusions of SA (red) or SE (blue) eEF1A2 proteins as a function of LifeAct-mCh levels as in panel E. Corresponding linear regression lines are also shown. (H) Coefficient of variation of LifeAct-mCh fluorescence fluctuations in spine pixels above a threshold from hippocampal neurons cotransfected with GFP-fusions of SA (red) or SE (blue) eEF1A2 proteins as in panel E. Median ± Q values are also plotted. (I) Hippocampal neurons were transfected with plasmids expressing mSc-actin and SA or SE GFP-eEF1A2 proteins, and actin mobility in dendritic spines was analyzed by FRAP. Bar, 1 μm. (J) FRAP profiles from dendritic spines as in panel I. Mean values (n>25) and fitted lines are plotted. The result of a paired t-test is also indicated (**** p<0.0001). (K) Mobile fraction of mSc-actin in single dendritic spines as in panel I. Median ± Q values are also plotted. The result of a Mann-Whitney test (* p<0.05) is also shown.

As F-actin is the major cytoskeletal protein in dendritic spines, we next tested whether phosphorylation of eEF1A2 in domain III regulates eEF1A2 dynamics at the synapse. Hippocampal neurons at 14 DIV were co-transfected with GFP-eEF1A2 proteins and LifeAct-mCherry as a marker for F-actin and, one day after transfection, we obtained time-series images to be analyzed by fluctuation analysis methods (Digman and Gratton, 2012). Notably, the phosphomimetic SE mutant showed a clear increase in the coefficient of variation (CV) of the fluorescence intensity over time in single pixels of spines when compared to the SA mutant (Figure 3E and F). These results could be explained by a higher propensity to dimerize, which would decrease the number of mobile fluorescent particles and, hence, increase the amplitude of fluctuations in the focal volume. However, ruling out this possibility, the SE mutant showed a lower dimerization efficiency than the SA mutant (Figure 3–figure supplement 1A). As an alternative, the lower fluctuation dynamics shown by the SA mutant in dendritic spines could be due to its association with relatively immobile structures such as the actin cytoskeleton. If this assumption were true, we would expect the local concentration of F-actin negatively correlate with the level of fluctuations. Results from single pixel analysis showed a clear negative correlation between LifeAct-mCherry levels and the fluorescence CV of the GFP-eEF1A2 SA mutant (Figure 3G). Furthermore, the accumulation of F-actin also produced the accumulation of the SA mutant protein at a single-pixel level in spines (Figure 3–figure supplement 1B). These results were also observed in dendritic axes (Figure 3–figure supplement 1C-E), supporting the idea that eEF1A2 phosphorylation prevents its interaction with F-actin and increases its mobility in spines and dendrites. Fluorescence fluctuations analysis at a single pixel level showed an increase in the coefficient of variation of LifeAct-mCherry in SE mutant compared to SA mutant (Figure 3H), indicating that the phosphomimetic form of eEF1A2 intensifies actin dynamics. To further explore this possibility, we measured mScarlet-actin mobility by fluorescence recovery after photobleaching (FRAP) in dendritic spines of neurons cotransfected with GFP-eEF1A2 proteins. Neurons expressing the phosphomimetic mutant showed faster recovery of mScarlet-actin fluorescence after photobleaching (Figure 3I-J). Furthermore, we detected a reduction in the mobile fraction of mScarlet-actin with the SA mutant (Figure 3K). In all, these results support the idea that the phosphorylation state of eEF1A2 regulates its interaction with actin and modulates actin dynamics.

### The phosphomimetic eEF1A2 mutant cannot sustain protein synthesis in yeast cells

In order to investigate the role of phosphorylation in translation, we decided to use budding yeast cells as an amenable model for precise genetic intervention. Since budding yeast cells possess two identical eEF1A-encoding genes *(TEF1* and *TEF2),* we used a strain in which the chromosomal copy of *TEF1* was disrupted and *TEF2* expression was under the control of a regulatable promoter. However, only mammalian Ser358 is conserved as Ser356 in yeast. We therefore expressed wt, SA or SE versions of *TEF1* under endogenous promoter *in trans* using a centromeric vector. Cells with empty vector grew slowly under *TEF2*-inducing conditions, but did not grow under repression conditions (Figure 4A). As expected, the presence of a full wt *TEF1* copy *in trans* fully rescued these phenotypes.

**Figure 4.**
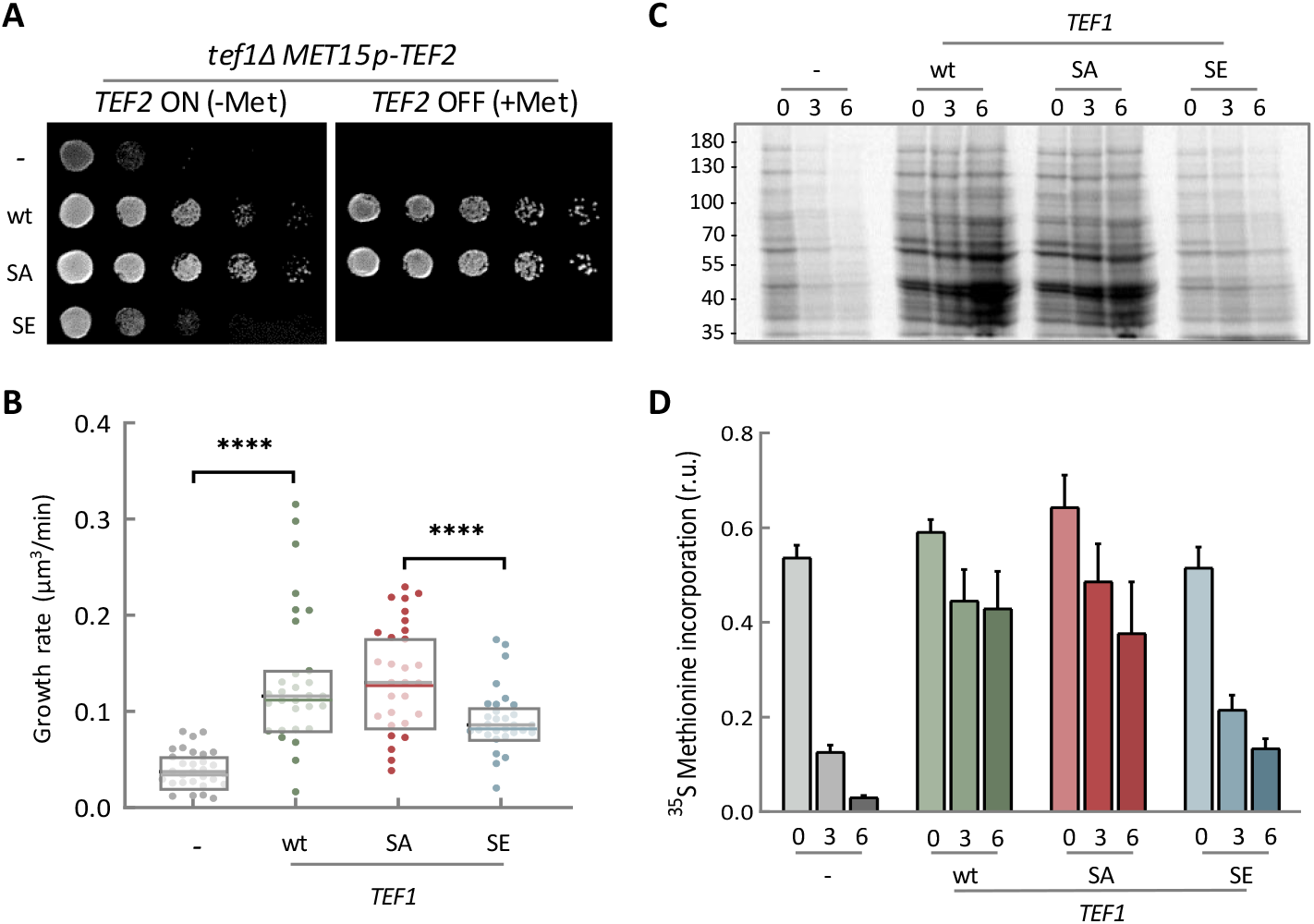
The phosphomimetic eEF1A mutant cannot sustain protein synthesis in yeast cells. (A) Growth of yeast cells expressing wt, SA and SE TEF1 proteins. Cells with a *tef1* deletion and *TEF2* expression under the control of *MET15p* as a methionine-regulatable promoter were transformed with centromeric vectors expressing wt, SA or SE versions of *TEF1* under endogenous expression sequences, and plated in the absence *(TEF2* ON) or presence *(TEF2* OFF) of methionine to test complementation by TEF1 proteins *in trans.* (B) Growth rate in G1 phase of cells as in panel A under *TEF2* repression conditions. Single-cell volume growth rates (n=30) and the corresponding median ± Q values are plotted. The results of Mann-Whitney tests are also shown (**** p<0.0001). (C) Pulse-labeling analysis of protein synthesis in cells expressing wt, SA or SE TEF1 proteins as in panel A. Cells were labeled for 5 min with ^35^S-methionine at 0, 3 and 6 h after *TEF2* repression, and lysates from equivalent culture volumes were analyzed by SDS-PAGE and autoradiography. Samples from control cells with empty vector (-) are also shown. (D) Quantification of protein synthesis rates analyzed as in panel C. Data are represented as mean ± SEM (n=3).

However, while the SA mutant was indistinguishable from wt, cells expressing the SE mutant were completely unable to grow under repression conditions. To ascertain defects in growth, we directly measured volume increase rates in G1 cells and observed the same results (Figure 4B). Finally, we performed pulse-labeling experiments 0, 3 or 6 h after promoter shut off (Figure 4C and D) and found that the phosphomimetic SE mutant caused a strong reduction in the overall protein synthesis rate. Therefore, these experiments support the notion that eEF1A2 phosphorylation at Ser358 inhibits protein synthesis in dendritic spines.

Considering the results related to the eEF1A2-actin interaction in neurons, we set out to investigate whether eEF1A2 phosphomutants would affect actin dynamics in yeast cells. Due to the lethal phenotype of SE mutant we were only able to analyze the SA mutant. Yeast actin cables assemble in the bud and bud neck and elongate into the mother cell during polarized growth from late G1 to the G2/M transition. To estimate actin cable growth, we photobleached Abp140p-GFP, an actin-binding protein, either at the bud neck (proximal bleaching) or at the opposite pole (distal bleaching, as control) and monitored loss of fluorescence in the middle third of the mother cell (Figure 4–figure supplement 1A). Since new actin monomers are added close to the bud neck, an increase in actin cable stability would favor Abp140p-GFP displacement and, hence, accelerate loss of fluorescence along the cable (Yang and Pon, 2002). We found that the SA mutant caused a significant increase in loss of fluorescence only if Abp140p-GFP was photobleached at a proximal position (Figure 4–figure supplement 1B), suggesting that the phosphoablated SA protein stabilizes actin cables in yeast cells.

### eEF1A2 stimulates translation in neuronal cells and interacts with its GEF in dendritic spines in a phosphosite-dependent manner

Given the severe translation defects caused by the phosphomimetic mutant in yeast cells, we decided to address this question in a neuronal cell line. To maximize the effect of transfected eEF1A2 proteins we created a stable Neuro-2a cell line expressing a shRNA against the 3’UTR of endogenous eEF1A2 and, after cotransfection of plasmids expressing GFP and eEF1A2 proteins, we carried out a puromycylation assay to visualize newly synthesized proteins (Figure 5A). We found that the SA mutant was able to stimulate translation (Figure 5B). In contrast, puromycin incorporation by the SE mutant was not significantly different from non-transfected cells. These results fully agree with the results obtained in yeast and confirm the relevance of eEF1A2 phosphosites in translation. Then, we decided to study the possible causes of this important functional output. Exchange of GDP for GTP is the first step in eEF1A2 recycling during translation, which is driven by eEF1B2 as catalytic component of the guanine nucleotide exchange (GEF) complex. To confirm the interactome analysis we decided to study the effects of phosphosites in the interaction between these two factors and performed immunoprecipitation analysis in HEK293T cells that had been co-transfected with HA-eEF1B2 and FLAG-eEF1A2 fusion proteins. We observed that the SE mutant was specifically affected, showing a 5-fold decrease in levels of co-immunoprecipitated eEF1B2 protein compared to wt (Figure 5C and D). By contrast, the phosphoablated SA mutant was as efficient as the wt eEF1A2 protein.

**Figure 5.**
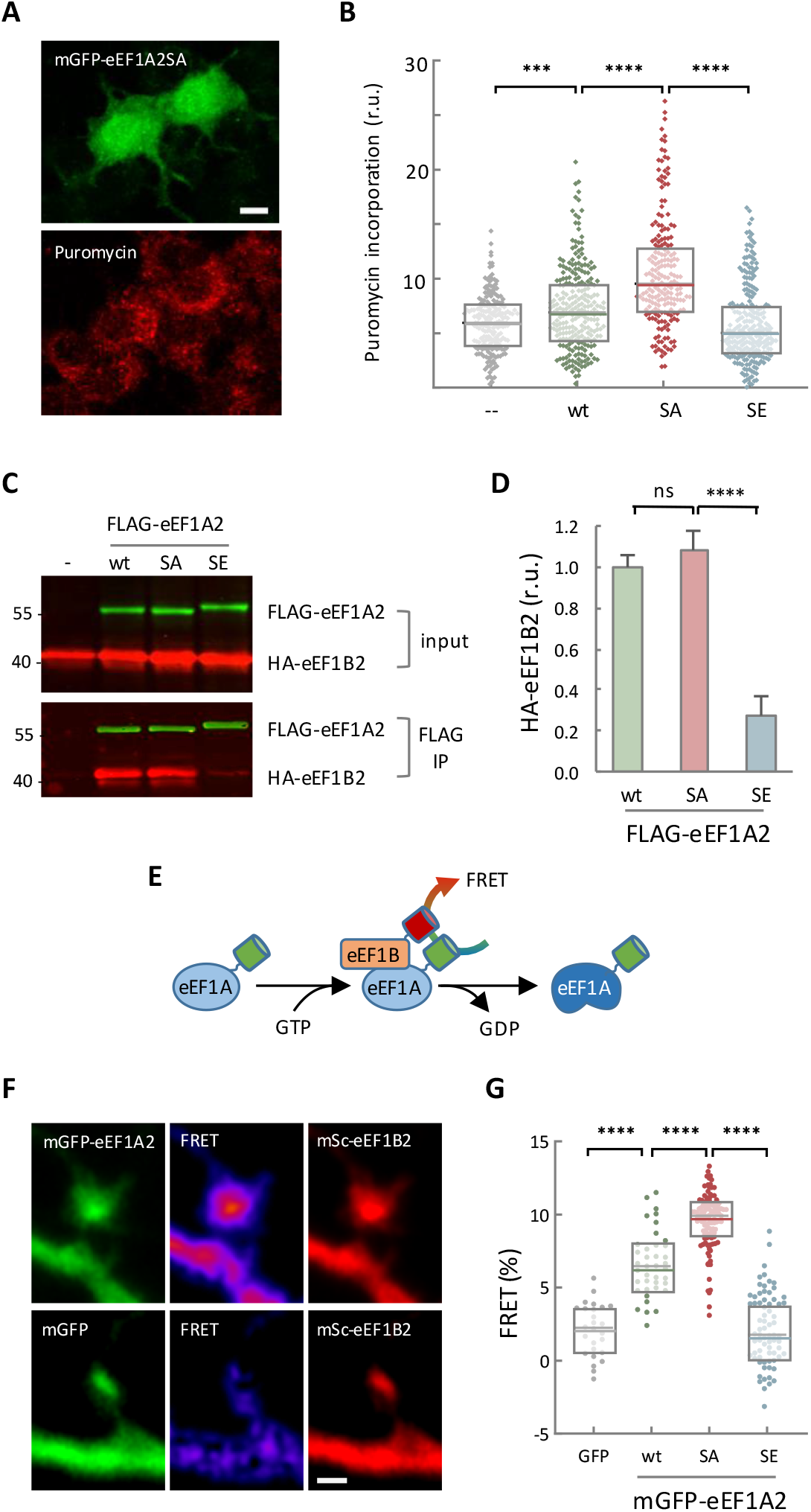
eEF1A2 stimulates translation and interacts with its GEF in dendritic spines in a phosphosite-dependent manner. (A) Neuro-2a cells stably expressing an shRNA against the endogenous eEF1A2 mRNA were cotransfected with plasmids expressing GFP and wt, SA or SE eEF1A2 proteins, and protein synthesis was assessed by puromycin incorporation and immunoblotting. Bar, 10 μm. (B) Quantification of puromycin incorporation in single Neuro-2a cells as in panel A. Median ± Q values (n>200) are also plotted and the results of Mann-Whitney tests (*** p<0.001; **** p<0.0001) are indicated. (C) Interactions between eEF1A2 and eEF1B2. HEK293T cells were cotransfected with plasmids expressing HA-eEF1B2 and FLAG-tagged wt, SA or SE eEF1A2 proteins. Immunoprecipitation was performed with αFLAG beads, and HA-eEFB2 and FLAG-tagged eEF1A2 protein levels in lysates (input) and immunoprecipitation (IP) samples were simultaneously analyzed by immunoblotting. (D) Quantification of eEF1B2 levels in IP samples from immunoblot analysis as in panel C. eEF1B2 protein levels were normalized relative to FLAG-tagged proteins in IP samples. Mean ± SEM (n=4), and the results of t tests (**** p<0.0001) are shown. (E) Schematic of the FRET strategy for quantifying the eEF1A2/ eEF1B2 interaction in dendritic spines. (F) Fluorescence and FRET images of representative spines from hippocampal neurons expressing mScarlet-eEF1B2 and mGFP-eEF1A2 or mGFP as control. Bar, 1 μm. (G) FRET levels in spines from hippocampal neurons as in panel F expressing mScarlet-eEF1B2 and mGFP or mGFP-tagged wt, SA or SE eEF1A2 proteins. The total number of observations (spines/neurons) analyzed is as follows: GFP, n=30/5; wt, n=40/6; SA, n=118/8; SE, n=75/7. Median ± Q values are plotted and the results of Mann-Whitney tests (* p<0.05; ** p<0.01; *** p<0.001) are also shown.

Next we wanted to visualize this interaction in dendritic spines where local translation plays an important role in synaptic plasticity. To this end, we measured Förster-resonance energy transfer (FRET) between mGFP-eEF1A2 and mScarlet-eEF1B2 (Figure 5E) and obtained interaction maps on spines from hippocampal neurons transfected at 13 DIV and imaged after 24 h. Notably, the SE mutant showed a significant reduction in FRET levels compared to the SA mutant and wt in spines (Figure 5F and G). Accordingly, the phosphoablated SA mutant showed the highest FRET levels, significantly higher than wt. Similar relative differences were obtained when FRET was analyzed in the soma of transfected Neuro-2a cells (Figure 5–figure supplement 1). In all, these results strongly support the idea that phosphorylation of the eEF1A2 factor could regulate its association with the eEF1B2 GEF and consequently modulate protein synthesis.

### DHPG induces transient phosphosite-mediated dissociation of eEF1A2 from its GEF in dendritic spines

Group 1 metabotropic glutamate receptors (mGluR1 and mGluR5) are implicated in different forms of mGluR-mediated synaptic plasticity that depend in part on regulation of local protein synthesis (Bear et al., 2004; Di Prisco et al., 2014; Muddashetty et al., 2007; Waung and Huber, 2009). Activation of mGluRs by DHPG stimulates the JNK pathway in cultured neurons (Yang et al., 2006) and has been linked to phosphorylation of key synaptic proteins such as PSD95 or elongation factor eEF2 (Nelson et al., 2013; Park et al., 2008). Moreover, polysome-associated JNK phosphorylates eEF1A2 at residues Ser205 and Ser358 as a response to DHPG in primary striatal neurons (Gandin et al., 2013). Taking all this data into consideration, we decided to analyze whether the activation of mGluRs with DHPG regulates the activity of eEF1A2 as an elongation factor. As expected, DHPG provoked a transient phosphorylation of eEF1A2 that reached its peak 4 min after treatment (Figure 6–figure supplement 1A and B). We then used the abovementioned FRET-based approach to analyze *in vivo* the association between eEF1A2 and its GEF eEF1B2. Remarkably, FRET levels in spines of neurons expressing wild-type eEF1A2 temporary dropped during the first 4 min after DHPG addition, indicating that DHPG causes a reversible reduction in eEF1A2-eEF1B2 interaction within a narrow time window after stimulation (Figure 6A and B). We noted that the fold-change reduction was stronger in spines with higher initial FRET values (Figure 6B and C). In sharp contrast, FRET levels produced by the phosphoablated SA mutant were maintained during DHPG treatment and did not correlate with the initial status of the spine. Thus, our results indicate that DHPG transiently downregulates the interaction between eEF1A2 and eEF1B2, thereby affecting the first step in the eEF1A activation cycle for translation elongation. Since the phosphoablated mutant was totally unaffected, the observed modulation would link activation of mGluRs, eEF1A2 phosphorylation and local modulation of translation in spines.

**Figure 6.**
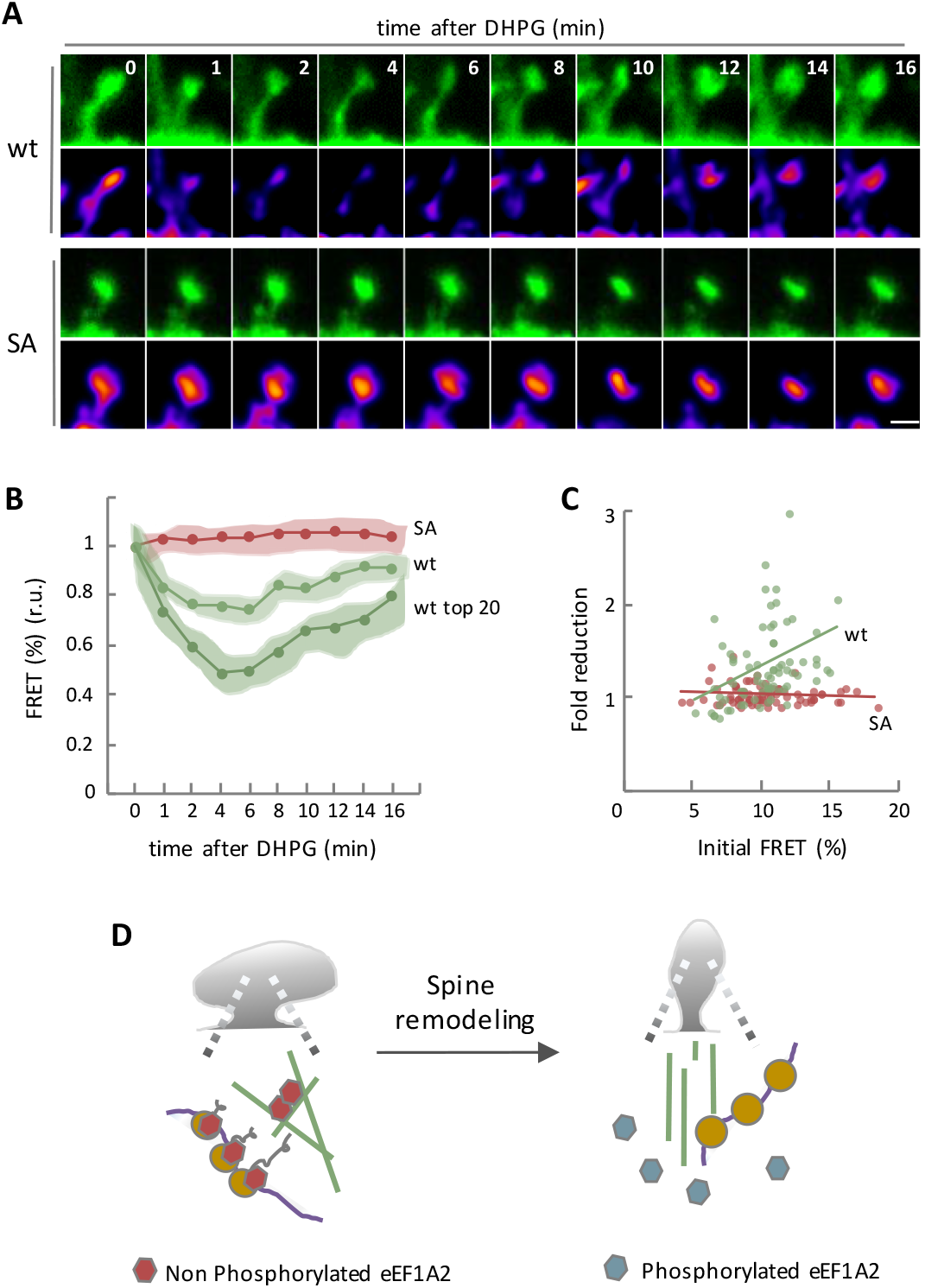
DHPG induces transient phosphosite-mediated dissociation of eEF1A2 from its GEF factor in dendritic spines. (A) Fluorescence and FRET images of representative spines from hippocampal neurons expressing mScarlet-eEF1B2 and mGFP-tagged wt or SA eEF1A2 proteins at the indicated times after DHPG addition. Bar, 1 μm. (B) FRET levels in spines from hippocampal neurons as in panel A expressing mScarlet-eEF1B2 and mGFP-tagged wt or SA eEF1A2 proteins as a function of time after DHPG addition. The total number of observations (spines/neurons) analyzed is as follows: wt, n=76/10; SA, n=131/12. FRET levels were made relative to time 0 and mean ± CL (α=0.05) values are plotted. Data from 20 spines with highest initial FRET values produced by wt eEF1A2 are also shown. (C) Transient fold-reduction in FRET as a function of initial FRET levels for mGFP-tagged wt (green) and SA (red) eEF1A2 proteins analyzed as in panel B. (D) Proposed role of eEF1A2 in dendritic spine remodeling. Two subpopulations of non-phosphorylated eEF1A2 would exist in stable spines, one involved in translation, likely in a monomeric conformation, and the other participating as dimers in F-actin bundles. As a result of synaptic stimulation, phosphorylation would dissociate eEF1A2 from F-actin and facilitate remodeling of the spine cytoskeleton. At the same time, eEF1A2 phosphorylation would cause its inactivation as a translation elongation factor, thus transiently preventing undesired protein accumulation before superimposed signals establish longer-term decisions as varied as LTP or LTD.

## Discussion

In this study we aimed to understand the physiological relevance of the eEF1A2 isoform in the context of synaptic plasticity. To this end, we focused our attention on Ser358 and three additional potential phosphorylation sites only present in isoform eEF1A2. Briefly, a phosphomimetic eEF1A2 SE mutant was seriously compromised in its ability to bind actin and produce actin bundles. Similar to the proposed role of CaMKIIβ in actin dynamics during LTP (Kim et al., 2015), dissociation of eEF1A2 would allow actin reorganization and activation of regulatory proteins related with actin cytoskeleton remodeling. Supporting this notion, our proteomic analysis showed a clear enrichment of actin binding proteins in SE immunoprecipitates. Actin binding proteins play roles in many different aspects of actin dynamics: polymerization, depolymerization, nucleation, branching, capping, cross-linking and trafficking (Pollard, 2016). Thus, according to previous studies in yeast (Perez and Kinzy, 2014; Umikawa et al., 1998), it is possible that eEF1A acts as a bridge between the cytoskeleton and actin modulators. Notably, as one of the proteins enriched in SE immunoprecipitates we found α-actinin-4, a Ca^2+^-sensitive actin-binding protein that interacts with group 1 mGluRs and orchestrates spine morphogenesis in primary neurons (Kalinowska et al., 2015). These data, along with the fact that the phosphoablated SA mutant causes a strong reduction in spine density in hippocampal neurons, point to the notion that phosphorylation of eEF1A is important for long-term structural plasticity.

In agreement with a role in structural plasticity, the eEF1A2 isoform has been implicated in metastasis (Abbas et al., 2015; Scaggiante et al., 2012; Tomlinson et al., 2005). It has been shown that eEF1A2 overexpression stimulates actin remodeling, cell invasion and migration (Amiri et al., 2007). A previous study in adenocarcinoma cell lines showed that eEF1A from metastatic cells has reduced F-actin affinity (Edmonds et al., 1996). Notably, eEF1A2 was found to be more enriched than eEF1A1 in cell protrusions of breast cancer cells (Mardakheh et al., 2015). These findings allow us to propose that localized eEF1A2 phosphorylation weakens its association with actin, increasing cytoskeleton reorganization, cell motility and finally, metastatic growth.

Synaptic activity has been reported to regulate the local translational machinery through changes in the phosphorylation status of initiation and elongation factors (Sossin and Costa-Mattioli, 2019). However, *in vivo* evidence for mechanisms regulating translation at a local level are still missing. Our FRET analysis to visualize the interaction between eEF1A2 and its GEF, the most upstream step in the translation elongation cycle, is the first direct evidence of a locally-modulated translation event in synaptic spines. Activation of mGluRs by DHPG stimulates the JNK pathway (Yang et al., 2006), and polysome-associated JNK phosphorylates eEF1A2 at Ser358 as a response to DHPG in primary striatal neurons (Gandin et al., 2013). Thus, our data on the behavior of phosphoablated and phosphomimetic mutants point to the notion that phosphorylation of eEF1A2 by JNK and/or other protein kinases mediating synaptic signals plays a key role in regulating local translation in dendritic spines.

Although local effects have not been demonstrated yet *in vivo,* a similar scenario has been described for eEF2 and translational suppression in cultured neurons (Marin et al., 1997; Park et al., 2008; Sutton et al., 2007), synaptic biochemical fractions (Scheetz et al., 2000) and hippocampal slices (Chotiner et al., 2003) after synaptic stimulation. This raises the question of whether inhibition of protein synthesis by the two elongation factors eEF1A2 and eEF2 are redundant mechanisms. Since both have been observed under similar mGluR stimulation conditions, phosphorylation of these two factors could be modulated by specific secondary signals. However, there is growing evidence that eEF1A also has a profound impact at the initiation step of protein synthesis. In yeast, mutations in eEF1A that affect aminoacyl-tRNA binding simultaneously cause actin binding and/or bundling defects but, intriguingly, increase phosphorylation of eIF2A by GCN2, the eIF2A kinase (Gross and Kinzy, 2007; Perez and Kinzy, 2014). Phosphorylation at Ser51 (conserved from yeast to mammals) by GCN2 converts eIF2A into an inhibitor of its own GEF eIF2B, leading to attenuation of general protein synthesis (Sonenberg and Hinnebusch, 2009). Therefore, regulation of eEF1A2 would offer at least two significant advantages compared to the eEF2 factor. First, modulation of GTP loading by eEF1A2 phosphorylation provides a mechanism to regulate the most upstream step in translation elongation. Second, phosphorylation of eEF1A2 could provide feedback on translation initiation and downregulate protein synthesis in a more efficient manner. Moreover, silent mRNAs would prevent subsequent initiation rounds and remain as monosomes as recently shown (Biever et al., 2020; Heyer and Moore, 2016).

The functional relevance of conserved Ser358 in protein synthesis is supported by our yeast experiments in which the phosphomimetic mutant showed a strong reduction in translation rates. We speculate that this phosphorylation event could be a mechanism for adapting yeast cells to specific situations. In this regard, it has been reported that glucose starvation causes rapid actin depolarization and inhibition of translation (Uesono et al., 2004). It remains to be determined whether phosphorylation of eEF1A2 plays any roles in this concurrent regulation of translation and actin cytoskeleton.

Our results shed some light on the purpose of the developmental switch between the two eEF1A isoforms. The transition of eEF1A1 to eEF1A2 is associated with development of the nervous and muscular systems (Lee et al., 1995). Downregulation of eEF1A1 during differentiation of these tissues is a general characteristic in vertebrates and yet is controlled through completely different species-dependent mechanisms, perhaps to establish specific mechanisms coordinating protein synthesis and cytoskeletal remodeling in terminally differentiated neurons, myocytes and cardiomyocytes (Newbery et al., 2007). According to these ideas, one of the aspects shared by these cells is a requirement for local structural plasticity. It has been shown that actin filaments have a role in maintenance of t-tubules in membranes of cardiomyocytes and myocytes (Vlahovich et al., 2009). The t-tubular membrane microfolds facilitate ion channel trafficking and modulate local ionic concentrations. Emerging evidence indicates that these microfolds generate very dynamic microdomains to modulate calcium-signaling processes (Hong et al., 2014). Furthermore, t-tubules are anchored to sarcomeric complexes whose maintenance depends on localized protein translation (Lewis et al., 2018). Thus, as we have found for dendritic spines, regulation of isoform eEF1A2 by phosphorylation could play a major role in microfold plasticity by regulating local translation and actin dynamics in sarcomeric Z discs, the t-tubule membrane-binding structure.

Our findings identify a novel mechanism by which metabotropic signaling regulates structural plasticity. The stimulation of mGluR increases Ca^2+^ levels, thus triggering activation of JNK and other Ca^2+^ signaling kinases (Giese and Mizuno, 2013). Here we show that receptor stimulation opens a time window in which elongation factor eEF1A2 dissociates from both its GEF protein and F-actin, thus decreasing protein synthesis and increasing actin cytoskeleton remodeling. This transitional state could be common to the different forms of synaptic plasticity including LTP, LTD and homeostatic plasticity, in which activity-dependent spine remodeling is an essential initial event (Figure 6D).

In summary, our work uncovers a crosstalk mechanism between local translation and actin dynamics in fast response to synaptic stimulation in neurons. As muscle cells also display a developmental eEF1A switch, we propose that eEF1A2 is a general effector of structural plasticity to attain long-term physical and physiological changes at the subcellular level.

## Materials and Methods

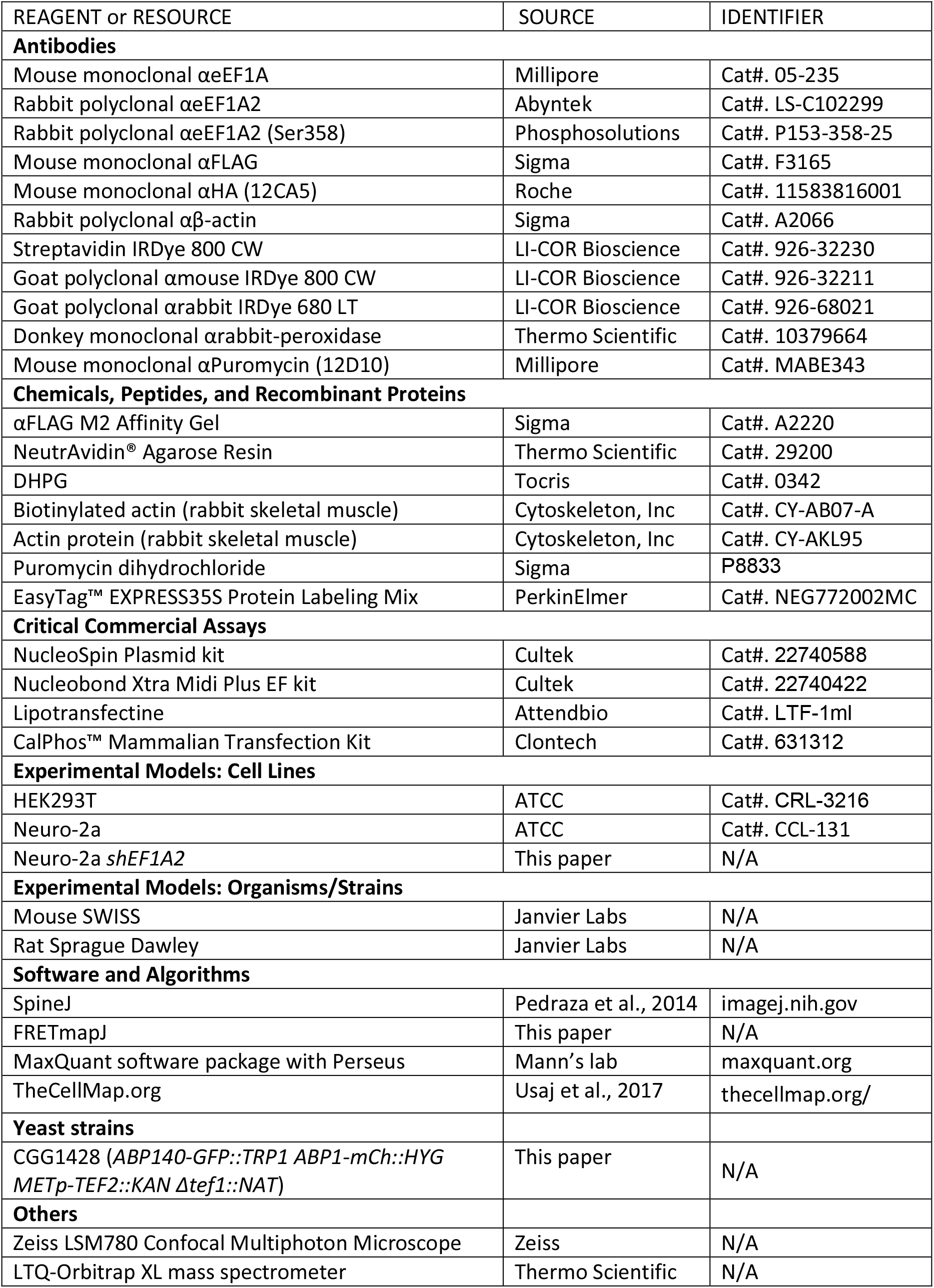
Key Resources Table

### Primary dissociated cultures

Animal experimental procedures were approved by the ethics committee of the Research Council of Spain (CSIC). Neurons were cultured as previously described (Pedraza et al., 2014). Briefly, hippocampi and cortex were isolated from E17 mouse embryos and digested with 0.05% trypsin at 37°C for 15 min. Dissociated cells were suspended in DMEM + 10% FBS + 0.6% glucose, and plated at a density of 5×10^4^ cells/cm^2^ for biochemistry and 2×10^4^ cells/cm^2^ for imaging experiments onto poly-D-lysine-coated plates. Medium was replaced 2 h after initial incubation with neurobasal medium (ThermoFisher, 21103049) supplemented with 2% B-27 supplement (ThermoFisher, 17504044), 1% GlutaMax (ThermoFisher, 35050061), and 1% penicillin/streptomycin. Neurons were placed in incubators at 37°C in 5% CO_2_. Medium was changed by half every 3 days.

### Hippocampal slice culture

Hippocampal organotypic slice cultures were prepared from postnatal day 6-7 rats as described (Bosch et al., 2014).

### Cell lines

HEK293T and Neuro-2a cells were grown in DMEM (Biowest, L0104) supplemented with 10% FBS (GE Healthcare Hyclone, 12350273).

### Yeast cells and cultures

Methods used for chromosomal gene transplacement were as previously described (Ferrezuelo et al., 2010). Cells were grown in SC medium with 2% glucose at 30°C without methionine to allow *MET2p-TEF2* expression, which was turned off by addition of methionine to 0.1 mg/ml when indicated.

### Gene transfection

Primary dissociated hippocampal neurons were transfected at 14 DIV using CalPhos mammalian transfection kit (Clontech) as previously described in Jiang and Chen, 2006 and analyzed 16h later. Organotypic slices cultures were biolistically transfected (BioRad) at 5-7 DIV and imaged 3-5 days later (Bosch et al., 2014). HEK239T and Neuro-2a were transfected using Lipotransfectine (AttendBio) according to the manufacturer’s protocol.

### DNA constructs

Site-directed mutagenesis in eEF1A cDNAs were performed by In-Fusion HD (Clontech), except domain III sequence containing the four mutations that was synthesized by GeneCustom. pcDNA3Flag5’, pcDNA3-6His-3HA, pEGFP-C3, pmScarlet-C1 and pET28A were used as host vectors. Plasmids were prepared using NucleoSpin Plasmid kit (Cultek) for cell line transfections and Nucleobond Xtra Midi Plus EF kit (Cultek) for neuron transfections.

### Real-time PCR analysis

Methods used for quantitative PCR have been described (Pedraza et al., 2014). Total RNA was isolated using E.Z.N.A. Total RNA Kit I (Omega Bio-tek) following the manufacturer’s instructions. All samples were treated with RNase-free DNase I (Thermo Scientific), and DNA contamination levels were assessed by qRT-PCR, omitting reverse transcriptase.

### Immunoblots and immunoprecipitations

Western blot analysis (Pedraza et al., 2014) was carried out with antibodies αeEF1A (Millipore, 1:1000), αeEF1A2 (Abyntek, 1:1000), αeEF1A2 (Ser358) (Phosphosolutions, 1:500), αFLAG (Sigma, 1:1000), αHA (Roche, 1:500), IRDye 800 (Li-Cor 1:10000), IRDye 680 (Li-Cor 1:10000), and streptavidin (LI-COR Biosciences 1:1000). Cell lysates from HEK293T cells were immunoprecipitated with αFLAG-agarose (Sigma).

### Interaction with actin

Actin pull-down *in vivo.* HEK293T cells were transfected with FLAG-eEF1A2 constructs, 24h post-transfection cells were harvested in collection buffer (20 mM Tris-HCl (pH 7.5), 50 mM KCl, 2 mM MgCl2, 1 mM ATP, 1% triton, 0.2 Mm DTT, 2 mM EGTA, and cOmplete EDTA-free and PhosSTOP from Roche). Supernatants were incubated with biotinylated-actin (Cytoskeleton, Inc), previously prepared as indicated by the manufacturer’s. NeutrAvidin agarose resin was used to pulldown actin complexes.

Co-sedimentation assay *in vitro.* F-actin bundling assay was carried out with purified eEF1A2 proteins from *E.coli* and lowspeed centrifugation. Actin was polymerized in the presence of 50 mM KCl, 2 mM MgCl_2_ and 1 mM ATP according to the manufacturer’s protocol. Recombinant proteins were incubated with F-actin for 30 min at RT and centrifuged at 4000g for 5 min to separate unbundled and bundled F-actin. Proteins were separated on SDS-PAGE and stained with Coomassie Brilliant Blue.

### Mass spectrometry based interactomic analysis

HEK293T cells were transfected with FLAG-eEF1A2 SA and FLAG-eEF1A2 SE expressing plasmids, and triplicate samples were immunoprecipitated using αFLAG-agarose beads (Sigma). FLAG immunoprecipitates (~150 μg protein) were reduced with 100 mM DTT at 95°C for 10 min, before being subjected to trypsin digestion using the Filter Aided Sample Preparation (FASP) protocol (Hau et al., 2020). Peptides were analysed using a LTQ-Orbitrap XL mass spectrometer (Barts Cancer Institute, London). MaxQuant (version 1.6.3.3) software was used for database search and label-free quantification of mass spectrometry raw files. The search was performed against a FASTA file of the *Mus musculus* proteome, extracted from uniprot.org. All downstream data analysis was performed using Perseus (version 1.5.5.3).

### Yeast growth rate in G1

Volume growth of yeast cells in G1 phase was measured by time-lapse microscopy in 35-mm glass-bottom culture dishes (GWST-3522, WillCo) essentially as described (Ferrezuelo et al., 2010) using a fully-motorized Leica AF7000 microscope.

### Protein synthesis measurements by pulse labeling

Strain CGG1428 expressing wild-type and mutant forms of eEF1A were grown as liquid cultures (100 ml) in medium lacking methionine at 30°C to OD_600_.= 0.2. Unlabeled methionine was then added to 50 mM to repress endogenous *TEF2* expresson and 0, 3 and 6 hr later cells were labeled for 5 min with 1 mCi/ml ^35^S Protein Labeling Mix (PerkinElmer). Lysates from 25-ml culture triplicate samples were analyzed by SDS-PAGE and autoradiography.

### Puromycin incorporation

Neuro-2a cells stably expressing an shRNA against endogenous the eEF1A2 mRNA were cultured on glass coverslips and transfected with GFP co-expressed with HA-eEF1A2 phosphomutants. 24 hours after transfection cells were treated with 1 μg/ml puromycin (Sigma) during 5 minutes and fixed in 4% paraformaldehyde in PBS. Immunofluorescence was performed as previously described (Pedraza et al., 2014), using α-puromycin (Millipore, 1:250) as primary antibody. Images were acquired with a Zeiss LSM780 confocal microscope. Immunofluorescence quantification was performed using ImageJ (Wayne Rasband, NIH). Puromycin incorporation was determined by measuring fluorescent intensity over the whole cell in transfected cells.

### FLIP imaging

FLIP was used as a quantitative assay to determine the stability of actin cables in yeast cells at room temperature in a Zeiss LSM780 confocal microscope equipped with a 40×1.2NA waterimmersion objective. A small circular region of the cell, either at the bud neck or at the opposite pole, was repetitively photobleached at full laser power while the cell was imaged at low intensity every 0.5 s to record fluorescence loss. After background subtraction, fluorescence data from an unbleached medial cell region were made relative to the initial time point, and a bleaching rate index was calculated as the inverse of the fluorescence half-life obtained by fitting an exponential function.

### Fluorescence fluctuation analysis

Hippocampal neurons were transfected with plasmids expressing GFP-eEF1A2 and LifeAct-mCherry. Fluorescence fluctuations were analyzed by time-lapse photon-counting microscopy using a Zeiss LSM780 confocal microscope with a 40 x 1.3 NA oil-immersion objective. Imaged regions were 248 x 100 pixels, with a pixel width of 86 nm/pixel at 13.0 μs/pixel.

### FRAP imaging

14 DIV hippocampal neurons cultured on 35 mm glass-bottom dishes (Ibidi) at 1.4 × 10^5^ cells/dish were transfected with plasmids expressing mScarlet-actin and SA or SE GFP-eEF1A2 proteins and analyzed 24h later. Live imaging was performed using Zeiss LSM780 confocal microscope equipped with a 5% CO_2_, 37°C humidified chamber under a 40×1.2 water objective. Photo-bleaching was achieved with 3 continues scans at maximum laser (561nm) power after 3 baseline images. Images were taken in 1 s intervals during 1 min. Photobleaching during the pre- and post-bleaching stages was negligible. FRAP efficiency was calculated using ImageJ. ROIs were placed on individual (bleached) spinesand non-bleached dendritic sections as control. Intensity values for spines were background subtracted and normalized to the average of the three pre-bleaching frames. Data were fitted to a single-term exponential recovery model as described (Koulouras et al., 2018).

### FRET imaging

Hippocampal cultures were transfected at 14 DIV with FRET biosensor plasmids expressing mGFP-eEF1A2 proteins and pmScarlet-eEF1B2. Time-lapse images were conducted 16h post-transfection. For neuronal stimulation experiments, hippocampal cultures were stimulated with 50 μM DHPG and images to calculate FRET efficiency were recorded every 2 min during 15 min at 37°C in 5% CO_2_. Neurons were imaged using a Zeiss LSM780 confocal microscope with a 40 x 1.2 NA water-immersion objective. Images were 1024 x 1024 pixels, with a pixel width of 65 nm. Briefly, donor (mGFP-eEF1A2) proteins were excited at 488 nm, and its emission was measured at 490-532 nm (I_D_). Excitation of the acceptor (pmScarlet-eEF1B2) was at 561 nm, and emission was measured at 563-695 nm (I_A_). We also measured the total signal emitted at 563-695 when excited at 488 nm (I_F_) to obtain the FRET efficiency as F%= 100 * (I_F_ – k_D_*I_D_ – k_A_*I_A_) / I_A_, k_D_ and k_A_ correcting acceptor cross-excitation and donor bleed-through, respectively, with the aid of FRETmapJ, a plugin that also provides maps with the FRET signal as pixel value for local quantification.

### Statistical analysis

Number of samples is described in the figure legends. Single spine data is displayed as median and quartile (Q) values. Pairwise comparisons were performed with a Mann-Whitney U test; and the resulting p values are shown in the corresponding figure panels. DHPG stimulation FRET data recorded from single spine during stimulation are represented as the mean value of the population along time, while the shadowed area represents the 95% confidence limits of the mean. Protein levels by immunoblotting and mRNA levels by RT-PCR were determined in triplicate samples and mean ± SEM values are shown.

### Data and software availability

SpineJ (Pedraza et al., 2014), BudJ (Ferrezuelo et al., 2010) and FRETmapJ can be obtained as ImageJ (Wayne Rasband, NIH) plugins from ibmb.csic.es/groups/spatial-control-of-cell-cycle-entry. The global yeast genetic interaction network (Usaj et al., 2017) can be accessed at CellMap.org.

## Acknowledgements

We thank M. Kerexeta and L. Pérez for technical assistance, C. Caelles for generous providing plasmids and proteins, I. Fita and M. Aldea for helpful comments and C. Rose for editing the manuscript. We also thank members of M. Aldea and C. Gallego laboratories for stimulating discussions and ideas. D.F.M. received a FI fellowship from Generalitat de Catalunya. This work was funded by grants from the Ministry of Economy and Competitiveness of Spain and the European Union (FEDER) (BFU2017-83375-R) to C.G.

## Author contributions

M.B.M., S.G., R.O., and C.G. built genetic constructs, primary cell cultures and performed the experiments. D.F.M. designed and performed yeast experiments. M.D., M.D. and F.K.M. contributed to mass spectrometry experiments. E.R. contributed to imaging experiments. M.B. carried out organotypic cultures. M.B.M., S.G. and C.G. conceived the study. C.G. analyzed the data and wrote the manuscript.

---

**Figure 1–figure supplement 1.**
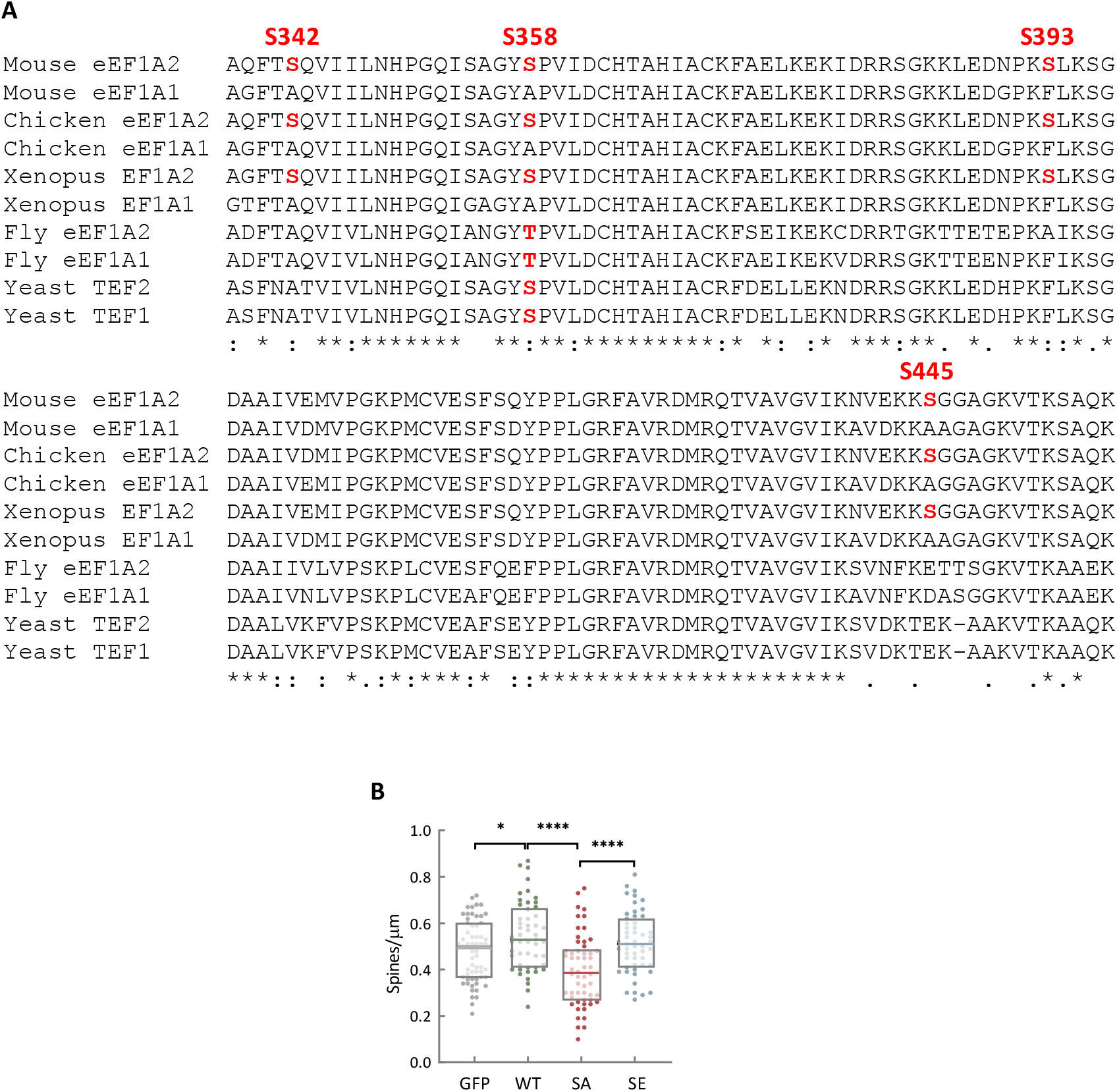
eEF1A2 phosphosite configuration modulates spine growth. (A) Sequence alignment of the eEF1A2 C-terminal domain from different species. Conserved serine residues mutated in this study are highlighted in red. (B) Quantification of spines per μm in hippocampal neurons transfected with plasmids expressing GFP or GFP-tagged wt, SA or SE eEF1A2 proteins. The total number of observations (spines/neurons) plotted is as follows: GFP, n=1990/61; wt, n=2040/51; SA, n=1434/56; SE, n=1952/53. Single-neuron data (dots) from three independent experiments and median ± Q values are plotted. The results of Mann-Whitney tests (* p<0.02; **** p<0.0001) are also indicated.

**Figure 2–figure supplement 1.**
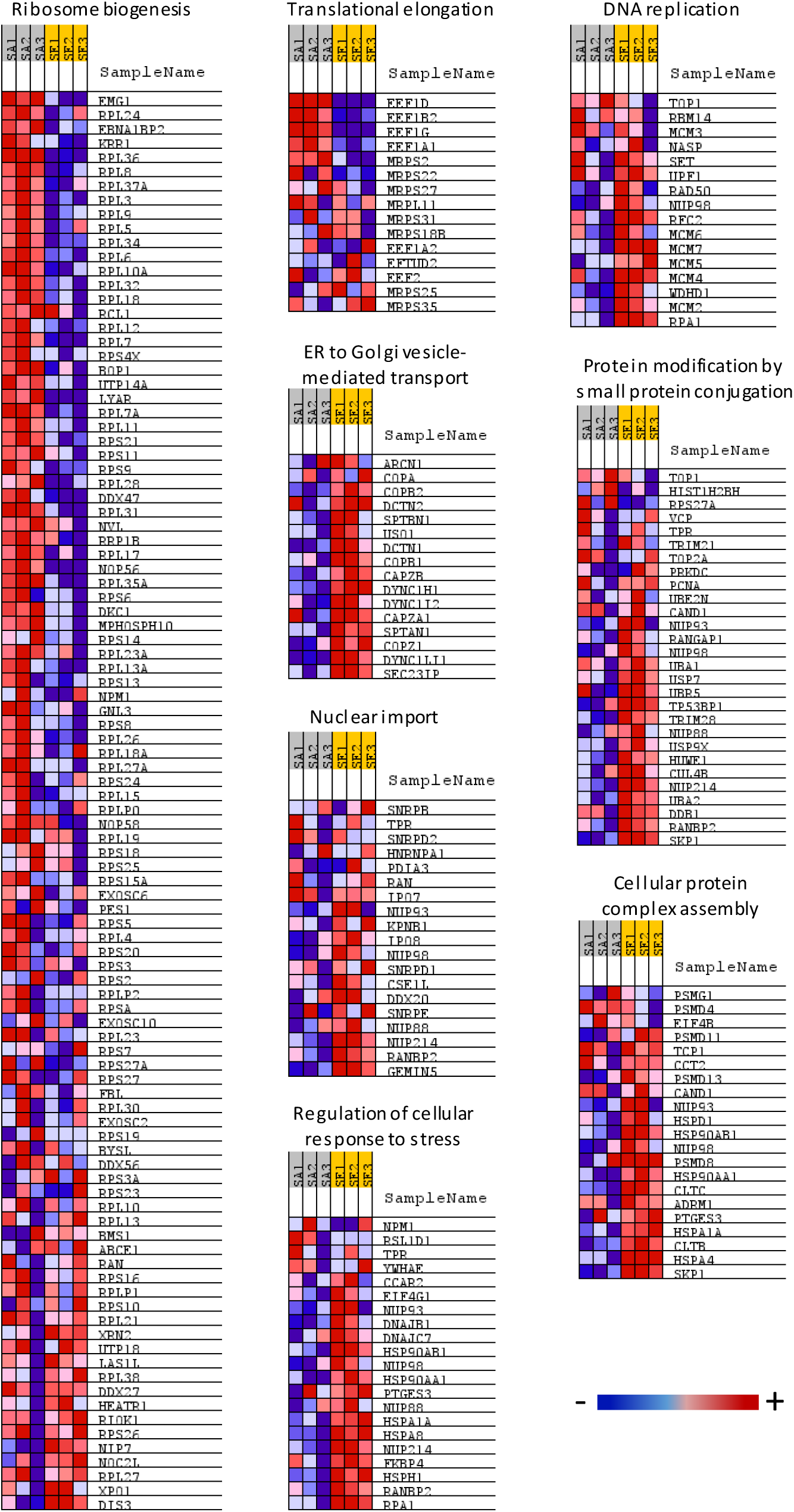
Interatomic analysis of eEF1A2 phosphomutants dissects translational and non-canonical functions. Triplicate immunoprecipitates from HEK293T cells expressing FLAG-tagged SA and SE eEF1A2 proteins were analyzed by LC-MS/MS. Colors in the heatmap denote high (red) to low (blue) normalized enrichment scores of individual proteins in the corresponding GO terms.

**Figure 3–figure supplement 1.**
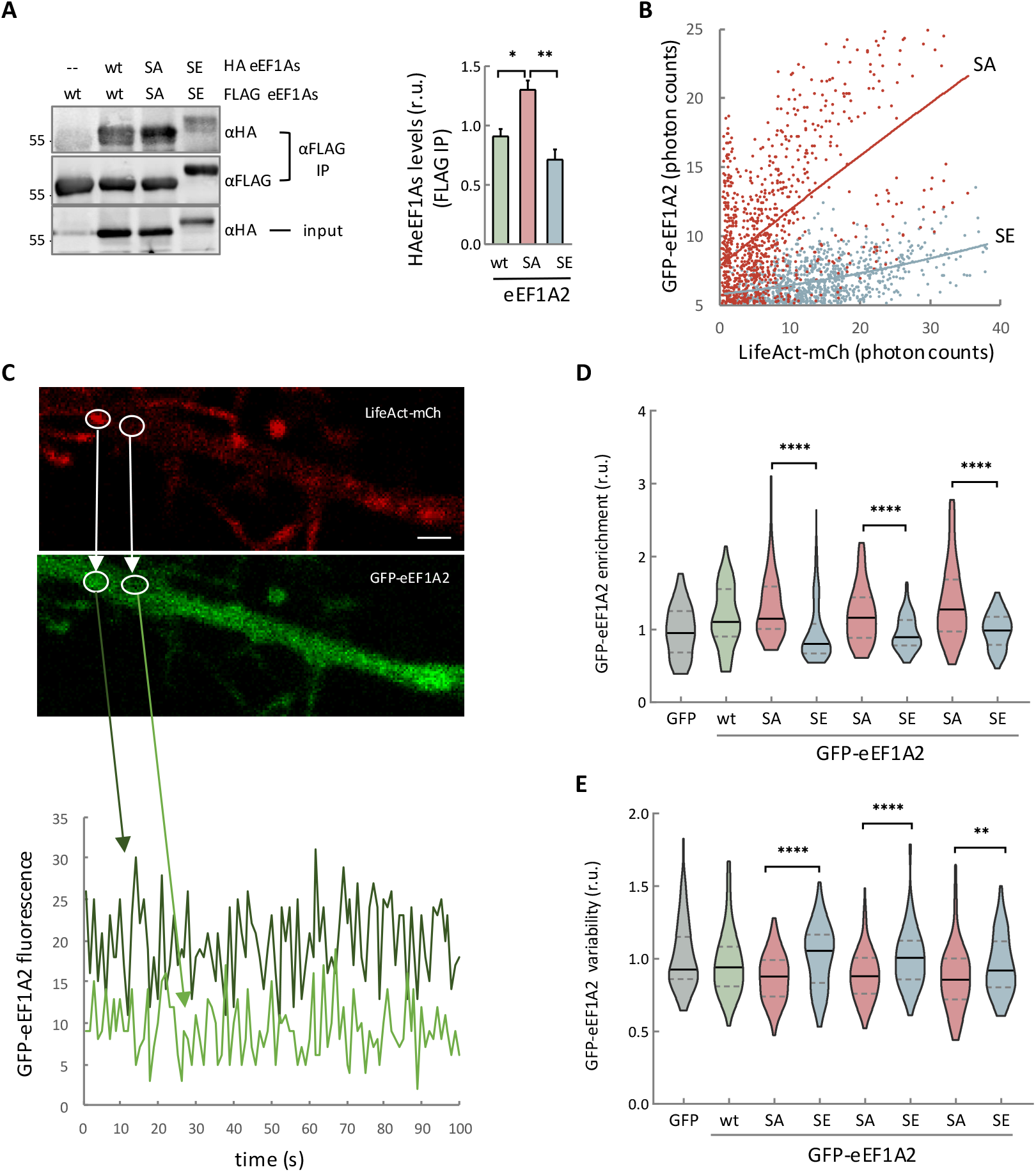
Phosphomimetic residues in eEF1A2 hinder its association with F-actin. (A) Immunoprecipitates (FLAG-IP) and lysates (input) from HEK293T cells expressing wt, SA and SE eEF1A2 proteins tagged with HA or FLAG as indicated were analyzed by immunoblotting. HA-tagged protein levels were normalized relative to FLAG-tagged protein levels in IP samples. Data are represented as mean ± SEM (n=4), and the results of t tests (* p<0.02, ** p<0.01) are also shown. (B) Hippocampal neurons were transfected as in Figure 3E and levels of GFP-fusions of SA (red) or SE (blue) eEF1A2 proteins as a function of LifeAct-mCh levels and linear regression lines are plotted. (C) Dendritic regions from hippocampal neurons as in Figure 3E were analyzed by time-lapse photon-counting microscopy. Representative temporal profiles obtained from GFP-tagged SA eEF1A2 in single dendritic pixels with high (dark green) or low (light green) LifeAct-mCh levels are shown at the bottom. Bar, 2 μm. (D) Levels of GFP-tagged wt (green), SA (red) or SE (blue) eEF1A2 proteins in dendritic pixels with high levels of LifeAct-mCh as in panel C. Cells expressing GFP (gray) are shown as control. Relative fluorescence levels in 200 pixels from 3 independent dendrites are plotted with median ± Q values for each condition. The results of Mann-Whitney tests (** p<0.01; **** p<0.0001) from triplicate samples are also indicated. (E) Coefficient of variation of fluorescence fluctuations from GFP-tagged wt (green), SA (red) or SE (blue) eEF1A2 proteins in dendritic pixels with high levels of LifeAct-mCh as in panel C. Cells expressing GFP (gray) are shown as control. Coefficients of variation in 200 pixels from 3 independent dendrites are plotted with median ± Q values for each condition. Results from triplicate experiments are shown for SA and SE mutant proteins. The results of Mann-Whitney tests (** p<0.01; **** p<0.0001) are also indicated.

**Figure 4–figure supplement 1.**
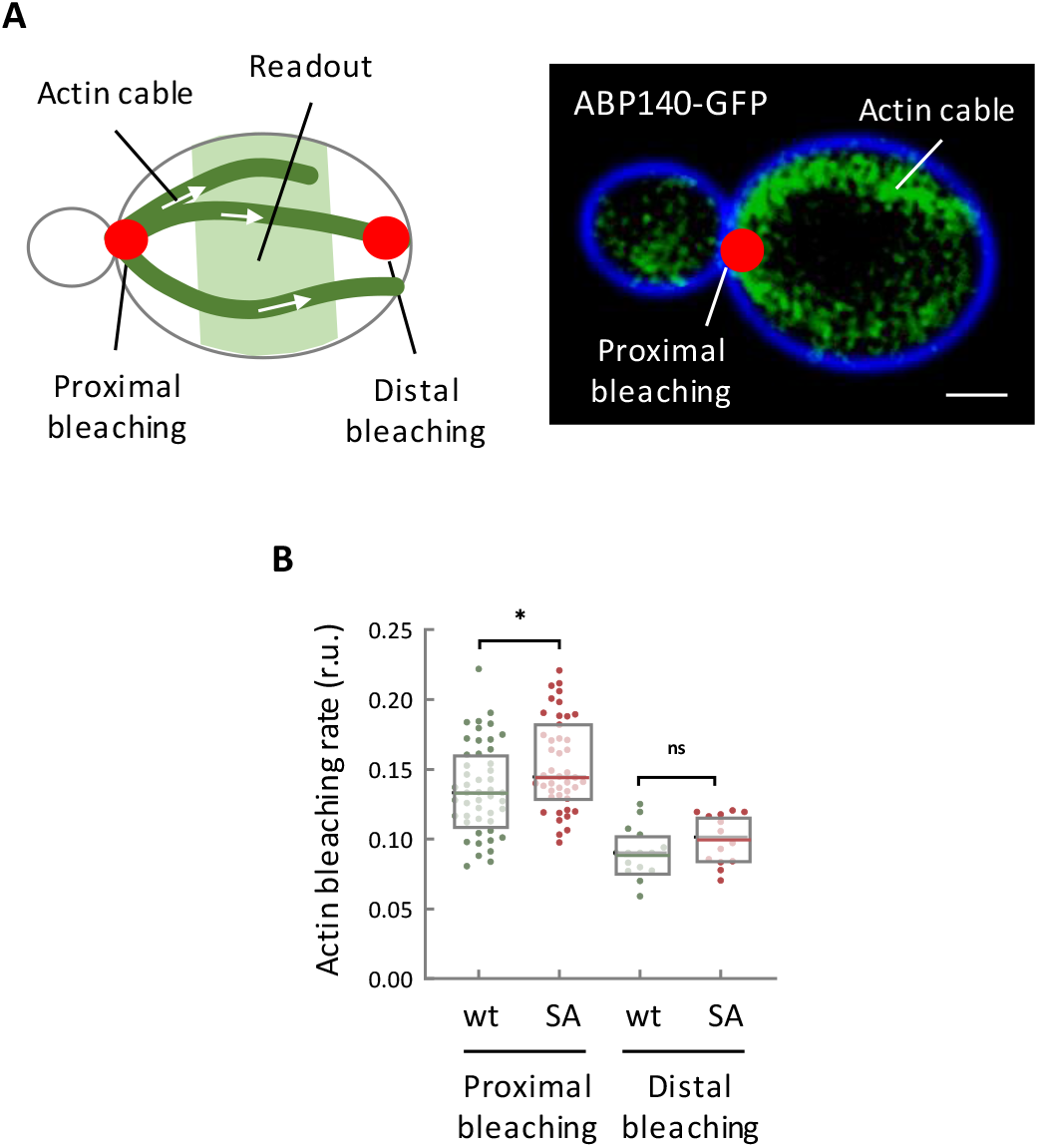
The phosphoablated eEF1A2 mutant promotes actin cable stability in yeast cells. (A) Schematic of a yeast cell showing the direction of actin cable growth from the bud towards the opposite pole of the mother cell (left panel). A representative image of actin cables as evidenced by Abp140p–GFP is also shown (right panel; bar, 1 μm). Due to actin cable growth, proximal bleaching has a stronger effect on FLIP readout at the middle third of the cell compared to distal beaching. (B) FLIP efficiency of Abp140p–GFP at proximal and distal positions in yeast cells expressing wt or SA TEF1 proteins under *TEF2* repression conditions as in Figure 4A. The number of observations (cells) analyzed is as follows: proximal bleaching wt, n=48; SA, n=44; and distal bleaching wt, n=15; SA, n=14. The results of a Mann-Whitney tests (* p<0.02; ns, non-significant) are also shown.

**Figure 5–figure supplement 1.**
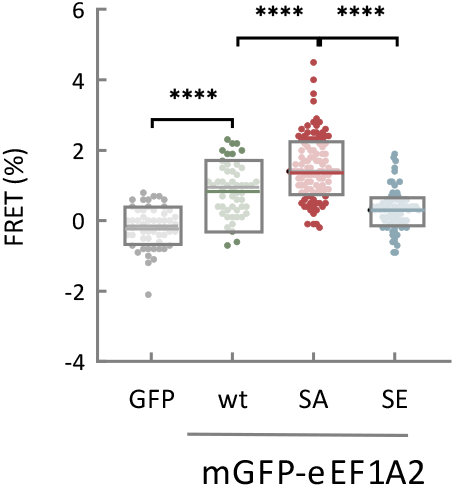
eEF1A2 interacts with its GEF in Neuro-2a cells in a phosphosite-dependent manner. FRET levels in Neuro-2a cells expressing mScarlet-eEF1B2 and mGFP or mGFP-tagged wt, SA or SE eEF1A2 proteins. The number of observations (cells) analyzed is as follows: GFP n=55; wt, n=53; SA, n=60; SE, n=48; all from three independent experiments. Median ± Q values are plotted and the results of Mann-Whitney tests (**** p<0.0001) are also shown.

**Figure 6–figure supplement 1.**
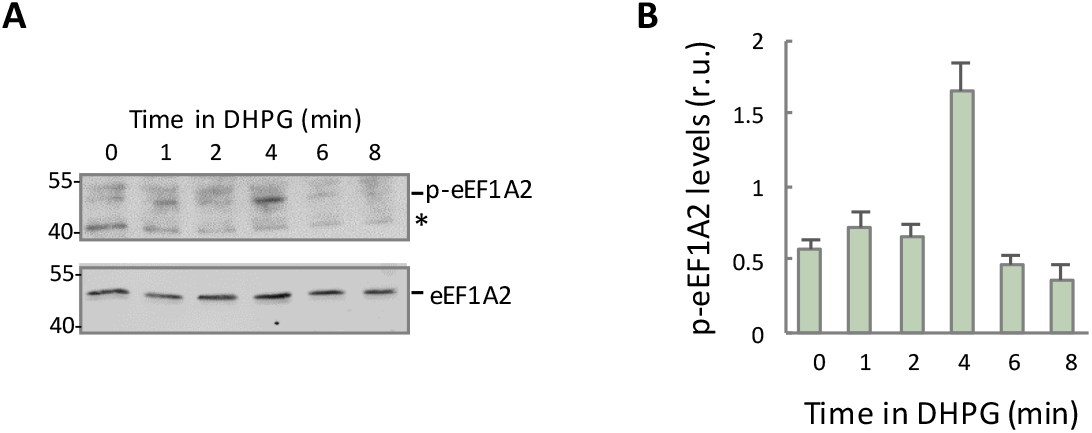
DHPG transiently phosphorylates eEF1A2 at Ser358. (A) Cortical cultured neurons were treated with DHPG and lysates were obtained at different time points for immunoblot analysis with p-eEF1A2 (upper panel) or total eEF1A2 (lower panel) antibodies. Asterisk indicates a nonspecific band. (B) Quantification of phospho-eEF1A2 from immunoblot analysis as in panel A. Phospho-eEF1A2 levels were normalized relative to eEF1A2 protein levels. Data are represented as mean ± SEM (n=4).

